# *Caenorhabditis elegans* model for *in vivo* screening of human gut microbiota mediated colonization and colonization resistance against *Clostridioides difficile*

**DOI:** 10.1101/2024.02.27.582212

**Authors:** Achuthan Ambat, Vishnu Thayyil Valappil, Sudeep Ghimire, Viju Vijayan Pillai, Phurt Harnvoravongchai, Shalabh Mishra, Sushim Gupta, Akhilesh Ramachandran, Purna Kashyap, Joy Scaria

## Abstract

The conventional bottom-up approach to probing the human gut microbiome’s link with hosts in germ-free models is hampered by considerable costs and time. To address this, our study introduces the nematode *Caenorhabditis elegans* as an innovative high-throughput model for exploring the gut microbiome’s impact on functional phenotypes. Traditionally, *C. elegans* studies have used continuous feeding for bacterial administration, a method that is unsuitable for anaerobes. For the first time, we have standardized a protocol for colonizing *C. elegans* with human gut anaerobes. By screening a microbial culturomics library representing 70% of the gut microbiome’s functional capacity, we showed successful colonization for 46% of the library. Functional phenotyping revealed that 5 of 10 strains, previously identified *in vitro* as inhibiting *C. difficile*, also inhibited *in vivo*. Validation of a selected strain in a germ-free mouse model confirmed colonization resistance and an immune response consistent with findings in *C. elegans*, underscoring the model’s translational potential.

## Introduction

A healthy, balanced human gut microbiota has profound effects on host immunity, metabolism, and resistance to infection^1–4^. Imbalances are increasingly implicated in conditions ranging from inflammatory bowel disease and recurrent *Clostridioides difficile* infection to obesity, type 2 diabetes, colorectal cancer, and neurodevelopmental disorders^5–8^. These effects depend not only on which microbial species are present, but also on how they interact. Disruptions in either dimension are often linked to pathological states^9,10^. To investigate these connections, microbiome studies have traditionally used a top-down strategy, sequencing and profiling whole fecal communities. While this approach maps diversity and links taxa to health or disease, it rarely resolves causal mechanisms^11–13^. For mechanistic insight, a complementary bottom-up strategy is needed, in which defined bacterial associations are tested under controlled conditions in a stepwise manner with increasing complexity. Such bottom-up strategies, involving mono-colonization or controlled combinations of species, can directly reveal how microbes influence host physiology^14^.

However, the vast diversity of the gut microbiota means that uncovering emergent interactions may require testing hundreds to millions of possible associations. Even with only 100 common gut species, the combinatorial space quickly becomes unmanageable: 100 mono-colonizations, nearly 5,000 pairs, over 160,000 triples, and more than 19 trillion possible communities when considering consortia of up to 10 members. Exhaustively testing these possibilities is impossible in practice. Rational design can narrow the space, but even then, hundreds of combinations may need to be tested. In higher animal models such as mice, this is feasible for tens of combinations but becomes too slow and prohibitively expensive at larger scales. This gap underscores the need for an intermediate system that bridges *in vitro* assays and rodent models to enable mechanistic testing of the gut microbiota at scale.

In this context, *Caenorhabditis elegans* offers a potential high-throughput model to fill this gap. It is a well-characterized model organism, introduced as a genetic system by Sydney Brenner in 1963, and remains widely used in biology^15–17^. *C. elegans* has a simple intestine that performs digestion and nutrient absorption, carries out functions analogous to the liver, and plays a central role in pathogen infection, immunity, longevity, and detoxification^18–22^. The culture and maintenance of this nematode are straightforward and economical. More recently, studies have shown that *C. elegans* itself harbors a microbiome, with around 300 bacterial genera identified, and community composition varying depending on environmental substrates. *C. elegans* has also been used to study interactions within soil microbiota, its natural habitat^23–25^. It has served as a host to test synthetic soil microbial communities, enabling mechanistic studies of simple to complex consortia and their emergent functions.

However, its application to human gut microbiota studies has been limited. The challenge arises because a large proportion of human gut microbes are obligate anaerobes, whereas both *C. elegans* and its native soil-associated microbiota are aerobic. This creates a paradox: gut anaerobes lose viability when introduced under aerobic conditions required for nematode survival, yet maintaining *C. elegans* under anaerobic conditions compromises worm viability. Resolving this incompatibility would unlock the potential of *C. elegans* as a high-throughput model system for studying human gut microbiota. In this study, we address this challenge by developing a standardized protocol that enables colonization of *C. elegans* with human gut anaerobes while maintaining worm viability, thereby opening the way for its use as a scalable platform for microbiome research. To implement this approach, we leveraged our well-characterized culture collection of gut microbiota from healthy human donors to test the feasibility of large-scale colonization in *C. elegans*^26^. Colonization was validated by fluorescence microscopy and CFU enumeration, confirming the reliability of the method. Analysis further revealed that mucin-associated sugar metabolism is a key determinant of successful colonization, supporting stable host–microbe interactions. Beyond single strains, we extended this framework to polymicrobial communities, showing that colonization resistance is shaped by community composition and that differential inhibition patterns reveal critical roles for specific taxa. These results demonstrate that the system can be applied not only for mechanistic studies of individual microbes but also for high-throughput screening of complex consortia.

Among the species tested, *Bifidobacterium longum* not only engrafted effectively but also conferred resistance to *Clostridioides difficile* infection, an effect reproduced in germ-free mice and highlighting the system’s translational potential. Analysis of immune responses in *C. elegans* further pointed to conserved pathways such as the p38 MAP kinase cascade, mirroring mechanisms described in mammalian hosts. Beyond individual strains, the model proved suitable for probing community-level interactions, revealing that colonization resistance is strongly influenced by composition and that critical taxa can be identified within defined consortia. Together, these findings establish *C. elegans* as a scalable *in vivo* platform for mechanistic studies of the human gut microbiota, enabling systematic screening of both individual microbes and complex communities, and bridging the gap between *in vitro* assays and mammalian models.

### Methodology

#### Experimental model and growth

##### Caenorhabditis elegans Strains

Mutant transgenic *C. elegans* strains used in this study are presented in the Key Resources Table. The Bristol strain N2 was used as the wild-type *C. elegans* strain (Brenner, 1974). All strains were maintained with an *E. coli* OP50 diet on Nematode Growth Media (NGM). All animals were maintained at 20°C unless otherwise indicated.

##### Mice Strains

We purchased six-week-old C57BL/6 Germ-free (GF) mice from Taconic Biosciences. Inc (New York, USA). To avoid any bias related to gender differences, we included both male and female mice in our experiments wherever it was possible. All animals were maintained in a 12-hour light and dark cycle with ad-libitum autoclaved drinking water and a standard chow diet. All animal experiments were performed with the prior approval of the protocols (19-014A, 19-044A) by the South Dakota State University (SDSU) animal studies committee and under the regulations of the Institutional Animal Care and Use Committee (IACUC) and SDSU policies.

##### Bacterial Strains

All the bacterial strains used in this study are presented in the Key Resources Table. All the anaerobic strains were grown in a Coy anaerobic chamber purged with 5% Hydrogen. All the media for anaerobic strain growth was purged with Nitrogen until the indicator (Resazurin) turns colorless.

### Method details

#### *C. elegans* growth conditions

Nematodes were handled according to standard practices (Brenner, 1974). Worm strains were grown on NGM plates for experiments with vegetative cells. All strains were grown at 20°C unless otherwise indicated. For regular strain maintenance OP50 at a concentration of 10x was used. For anaerobic bacterial colonization experiments, the same 10X concentration was used and seeded 0.5mL for 100mm plates and 0.05mL for 60mm plates and spread using an L-rod for forced feeding. For dead *E. coli* OP50, the pre-frozen OP50 pellet was heated at a temperature of 90 C in a water bath for 10 min, resuspended in M9 to a 10x concentration, and seeded 0.5mL for 100 and 0.05mL for 60mm plates, for more than three days of colonization or the life span assay NGM containing 50mM Fluroxydiuridine (FUDR) was used. For high through colonization, the worms were fed in anaerobic axenic media^1^ containing specific bacteria of interest, as mentioned, and then transferred to aerobic axenic media containing 10x dead *E. coli* OP50.

#### Bacterial growth conditions

All the anaerobic bacteria for the study were grown in modified Brain Heart Infusion (mBHI) broth, as mentioned previously in ^2^. Briefly, the media was prepared by adding yeast extract (5.0 g/L), L-cysteine (0.3 g/L), and 1 ml/l Resazurine (0.25 mg/ml) to standard BHI base ingredients. The media was purged until anaerobic and was autoclaved at 121°C and 15 lbs pressure for 30 minutes. The media was then transferred to the anaerobic chamber and further supplemented with 1 ml/L of hemin solution (0.5mg/ml), 0.1 ml/L Menadione (58 mM), 10 ml of ATCC vitamin mixture (ATCC, USA), and 10 ml of ATCC mineral mixture (ATCC, USA). For making *Clostridium difficile* selective agar, mBHI was supplemented with 0.3g/L of D-cycloserine and 0.002g/L of Cefoxitin. *Bifidobacterium* selective Medium was made as per the manufacturer’s instruction. Finally, *E. coli* OP50 was grown in Luria Bertani media aerobically unless mentioned otherwise.

#### *C. elegans* gut bacterial colonization and enumeration

Colonization of anaerobic and aerobic bacteria was done similarly. The axenic L1 worms were generated as previously mentioned^3^. First, 8-10 of 3-4 days-old NGM plates were washed with 5mL sterilized M9 buffer. Next, the worms were washed twice with 10mL of M9 buffer. Worms were then treated with 5mL bleach solution (Bleach:5N NaoH:Water at a ratio 2:1:2) and vigorously shaken for 3 minutes. Finally, bleached eggs were washed twice with 10mL of M9 buffer, redissolved in 2mL of M9, and incubated in a rotor shaker at room temperature for ∼10 hours.

Axenic worms were then anaerobically exposed to previously anaerobically seeded NGM plates with 500uL of 10x *B. longum*, *C. difficile* R20291, *C. difficile* 43598, or *E. coli* OP50 for 3 hours. After the exposure, plates were placed in aerobic condition for another 1 hour. The worms were transferred to a microcentrifuge tube with 1mL of M9 buffer, washed 3-4 times, and resuspended in ∼100uL of M9. The worms were then transferred to previously seeded NGM plates with dead OP50. For the colonization experiment of more than 3 days, worms were picked and transferred onto NGM plates containing FUDR.

For enumeration, worms were transferred from the plates using 5mL of M9 and washed at least 8-10 times with 10mL of M9. The worms were then exposed to 5mL of 5mg/mL of vancomycin in M9 for 40-50 minutes. Worms were then washed thrice with M9 and redissolved in 200uL of M9. Next, pulverization was done, as previously mentioned using a mortar and pestle ^4,5^. Pulverized worms were then diluted in PBS and plated on Bifidobacterium selective medium (BSM) for *B. longum*, *Clostridium difficile* selective agar (CDSA) for *C. difficile* and Luria Bertani agar for *E. coli OP50*.

#### High throughput colonization

10 mL of previously grown anaerobic bacteria as mentioned in (Table S1) was pelleted down and resuspended in 1 ml of anaerobic axenic media. In a 96-well 1mL block containing 800 µl of axenic media,100 µl of bacterial suspension and 100 µl of worms (∼1worm/uL) in anaerobic axenic media were added. The plate was sealed with a breathable membrane (Sigma) and incubated anaerobically for 3 hours at 25 . Following this, the plate was aerated for 30-45 minutes in a laminar hood. After that the worms were washed with 500 µl of aerobic axenic media for 3-4 times and then resuspended in 500 µl of axenic media. The worms were then treated with 500 µl of heat-killed OP50 in axenic media, the plate was resealed with breathable membarne, and incubated at 20 for 24 hours. For bacterial isolation, the worms underwent multiple washings using 0.1% Triton X in M9 buffer, including treatment with 0.2% bleach in M9 for 6 minutes at 4 . Finally, worms were disrupted with silicon grits, and the bacterial load was quantified by serial dilution and plating on mBHI agar, followed by incubation at 37 overnight. For each batch of the experiments, *B. longum* and *E. coli* OP50 served as a positive control, and no bacteria treatment served as a negative control.

#### Colonization resistance in *C. elegans*

##### Mono bacterial

For colonization resistance, initial colonization of *B. longum/C. difficile* or *E. coli OP50* was done per the previously mentioned colonization protocol. After 24 hours of incubation in NGM plates with dead *E. coli* OP50, the worms were washed and re-inoculated anaerobically with previously seeded NGM plates with *C. difficile/B. longum* or *E. coli* OP50 and incubated for 3 hours. After, the plates were placed in an aerobic condition for another 1 hour. Finally, the worms were transferred into microfuge tubes washed 3-4 times with M9 and resuspended in ∼100uL of M9. These worms were then transferred to NGM plates with dead OP50. Enumeration of *C. difficile* was done in CDSA, as previously mentioned.

##### Poly bacterial

For colonization resistance in poly bacterial associated worms, top 10 invitro inhibited bacteria (*Bifidobacterium bifidum, Faecalibacterium praunitzii, Anaerostipes hadrus, Eisenbergiella tayi, Lactobacillus muris, Bifidobacterium longum, Parabacteroides dorei, Bacteroidetes eggerthi, Bacteroidetes thetaiotaomicron,* and *Bacteroidetes adolescentis)* was grown in mBHI for 24 hours. Post 24 hours OD (600nm) was taken for each culture normalized to 0.5 mixed in equal volume. The culture was pelleted down and made 10x concentration and 500uL was seeded onto NGM plate. Further colonization was done as previously mentioned.

After 24 hours of incubation in NGM plates with dead *E. coli* OP50, the worms were washed and re-inoculated anaerobically with previously seeded NGM plates with *C. difficile* and incubated for 3 hours. After, the plates were placed in an aerobic condition for another 1 hour. Finally, the worms were transferred into microfuge tubes washed 3-4 times with M9 and resuspended in ∼100uL of M9. These worms were then transferred to NGM plates with dead OP50. Samples were also aliquoted for Microbiome sequencing. Enumeration of *C. difficile* was done in CDSA, as previously mentioned.

#### *In-vitro* coculture inhibition and antimicrobial inhibition

Coculture inhibition was performed as previously mentioned in ^2^. Briefly, for the antimicrobial inhibition assay, *B. longum* strains were grown for 24 hours in mBHI medium and then centrifuged at 10,000 g for 5 min. The supernatant was filter-sterilized using a 0.22 μ filter in the anaerobic chamber and diluted in a ratio 1:1 with the 1X mBHI medium. pH was adjusted to 6.8. An overnight culture of *C. difficile*R20291 was adjusted to OD600= 0.5. Twenty microliters of OD600 adjusted suspension was added to 1 ml of the 1:1 diluted cell-free supernatant and incubated for 24 hours in triplicate. *C. difficile*R20291 was grown in 50% mBHI diluted with anaerobic PBS as a positive control. After 24 hours, the cultures were serial diluted with anaerobic PBS, plated on CDSA, and incubated anaerobically for 24 hours for enumeration. For Heat treatment, the centrifuged supernatant was transferred to sterile tubes and pasteurized at 90 °C for 45 min (1 mL portions) suspended in a water-filled heating block. Pasteurized supernatant preparations were stored at 4 °C for near-term experiments or frozen at 20 °C. For Proteinase K, portions of supernatants were treated with proteinase K) at a concentration of 1mg mL-1 for 1 hour at 37°C. Following treatment, the enzyme was inactivated by the addition of phenylmethylsulfonyl fluoride.

#### Avoidance Behavior and death assay

Avoidance behavior was measured as mentioned previously ^6^ with slight modification. Briefly, NGM plates were anaerobically seeded with 5x concentration of *B. longum*, *C. difficile* R20291, *C. difficile* 43598, or *E. coli* OP50. The plates were then inoculated with ∼30 worms onto the center of the previously seeded NGM plates. The number of worms inside and outside the bacterial lawn was counted at different life stages, and percentage avoidance was calculated. Dead worms / Crawled out worms were not included in the analysis. To evaluate the toxicity of strains, the number of dead/alive worms was counted at different life stages. Life stages for worms were different for different bacteria.

#### Life span Analysis

Life Span analysis was performed as mentioned previously ^7^ with slight modifications. Briefly, colonization of *B. longum*, *C. difficile* R20291, *C. difficile* 43598, or *E. coli* OP50 was done as discussed earlier and transferred to NGM plates containing heat-killed *E. coli* OP50. During the L4 stage of their life cycle, the 20-30 worms were transferred to NGM plates supplemented with 50mM FUDR and seeded with heat-killed *E. coli* OP50. The worms were then observed using the dissecting scope daily for their viability, and data were recorded. Worms that crawled out of the plate were omitted from the analysis, and dead worms were removed. Worms are picked up and transferred to new NGM with FUDR plates weekly to replenish the food.

#### Mice experiments

For simple consortium gnotobiotic mice, *B. longum/C. difficile* was used for inoculation. For *B. longum* pre-treatment, mice were gavaged with a volume of 200uL of 20x concentrated *B. longum /* sterile saline freshly prepared for 2 days. The mice were undisturbed for the next 7 days until stable colonization was observed. For *C. difficile* pathogenesis, we used 200uL of 10^4 CFU of *C. difficile R20291* for consecutive 2 days, 7 days post *B. longum*/saline gavage. A volume of 200uL of the bacterial mix was orally administered to the treatment group. Mice were sacrificed 5 days post *C. difficile* infection. We used a triplicate treatment group with 4-6 mice in each group, while a group of uninoculated mice was used as GF control.

#### Bacterial enumeration

Total bacterial count in the fecal samples collected at different time points was assessed to evaluate the bacterial load, which indicates the bacteria’s colonization. Anaerobic mBHI/CDSA was used to enumerate the bacterial load used in this study.

#### RNA Isolation and RT-PCR

##### C. elegans

RNA isolation was performed using Trizol reagent, as previously mentioned in ^8^, with slight modifications. First, continuous colonized worms were harvested at 24-hour intervals, washed off from the plate using 5 mL of M9, and transferred into a centrifuge tube. The worms were washed at least five times at 1500 rpm 1min to remove any existing bacteria. Next, the worms were resuspended in 500uL of Trizol reagent and transferred to a microfuge tube. The tubes containing the worms were then snap frozen in liquid nitrogen and thawed at 37°C at least ten times for the worms to lyse properly. The Trizol-containing worms were then pipetted up and down until all the worms were sufficiently dissolved.

Next, 100uL of chloroform was added to the lysed Trizol reagent, vortexed properly, and incubated for 3 min at 25°C. The Trizol/chloroform mixture is centrifugated at 12500rpm for 15 min at 4°C. The aqueous phase was separated into a fresh tube, and 250uL of cold isopropanol was added. The mixture was vortexed and incubated for 10 min at 25°C. Centrifugation was done at 12500 rpm for 10 min at 4°C. The supernatant was discarded slowly by not disturbing the pellet at the bottom of the tube. The pellet was further washed with 500uL of 80% ethanol by centrifugation at 7500 rpm for 5 minutes at 4°C. The pellet was further dried in air for 3 min to remove the residual ethanol. The pellet was then resuspended in 20uL of nuclease free water, and the concentration was checked using an RNA HS assay kit in Qubit 4 fluorimeter. RNA was stored at -80°C until used.

50 ng of total RNA (see above) was reverse transcribed according to the manufacturer’s protocol (Protoscript II, NEB). The reaction was done in technical triplicates, and the product was pooled together and diluted with 80uL of NFW. Reverse transcribed cDNA was frozen at -20°C until further use. Q-PCR was done using Syber green PCR Master mix (Invitrogen) with a primer concentration of 10uM with 4uL of cDNA product. Q-PCR was done in Quantstudio Real-time system following the program; 50°C for 2 min, 95°C for 10 min, and 40 cycles of 95°C for 15 sec, 55°C for 30 sec. We used two housekeeping genes, Actin and Peroxisomal membrane protein-related genes, for calculating the fold change of gene expression relative to worms fed with *E.coli OP50* using the 2^-ΔΔCt^ method.

##### Mice colon

To assess the localized response of host-microbe interaction resulting from the colonization of *C. difficile* and *B. longum*, 100mg of tissue from the colon was sampled and snap-frozen in liquid nitrogen. The samples were then saved at -80°C until further processing. For RNA extraction, we used a TRIzol® (Ambion, life technologies)- chloroform (Sigma-Aldrich) method. According to the manufacturer’s protocol, a DNase treatment was performed using an RNase-Free DNase kit (Qiagen, Maryland) to remove any contaminant genomic DNA. The RNA quality and quantity were evaluated using NanoDrop One (Thermo Scientific) and saved at -80°C until further use.

Complimentary DNA (cDNA) was prepared from 250 ng of total RNA using the First strand cDNA synthesis kit (New England BioLabs Inc.) as per the manufacturer’s protocol. Host gene expressions were assessed by qRT-PCR using the 25 define immune-related genes. Reactions were prepared using Power SYBR® Green PCR Master mix (Applied Biosystems). PCR run was performed in an ABI7500 standard (Applied Biosystems) RT-PCR machine using the following cycling conditions: 95°C for 10 mins, 40 cycles of 95°C for 15 sec and 60°C for 1 min.

Raw cycle threshold (C_T_) values at a threshold of 0.15 were exported and then uploaded to the Qiagen data analysis center for further analysis. A C_T_ cut-off value of 30 was set for analysis. An average geometric mean of C_T_ values of two housekeeping genes - mouse beta-actin (Actb) was used for data normalization and ΔC_T_ calculation. Fold change and fold regulation in the gene expression were calculated using the ΔΔC_T_ method.

#### Microbiome DNA extraction and Sequencing

For DNA extraction from colonized microbiome member of *C. elegans* intestine. 30 -50 worms wer picked and put it in 500μL of M9. worms were then washed at least 5 time in M9 to remove any external bacteria. Post washing the worms are pulverized, as previously mentioned using a mortar and pestle ^4,5^. Genomic DNA was extracted from pulverized worms using a DNeasy power soil kit (Qiagen, Maryland, USA), according to the manufacturer’s instructions. The 16s rRNA gene sequencing was performed according to the standard Illumina protocol, where PCR amplicons targeting the V3-V4 region of the bacterial 16s rRNA gene were used for sequencing.

### Imaging

#### CFDA-SE staining

CFDA-SE (5,6-carboxyfluorescein diacetate succinimidyl ester) staining was done as per manufacturer’s protocol. For colonization, the NGM plates were seeded with stained *B. longum*, *C. difficile* R20291, *C. difficile* 43598, or *E. coli* OP50 3 hours before worm inoculation anaerobically. Worms were then inoculated anaerobically, and after 3 hours, the plates were kept in aerobic condition for another 1 hour. Visualization of bacteria was done in the 4^th^ hour. Agar pads were made on the microscopic slide using 2% agarose for fixing the worms. Then, 10uL of stained bacteria-treated worms were placed on the slides, and 5uL of 1mM levamisole was added to anesthetize the worms. The worms were covered with a coverslip and visualized under a fluorescent microscope with FITC filter illumination (419/516 Ex/Em).

#### Transgenic worms

Colonization and colonization resistance was done for the Transgenic worm, same as the wildtype mentioned before for MAH236, CF1139, and QQ202 *C. elegans* strains tagged with GFP. The worms were visualized 3, 24, and 48 hours after the colonization. Agar pads were made on microscopic slides using 2% agarose to fix the worms. 10uL of worms at 3 and 24 hours were placed on the slides, and for 48 hours, individual worms were picked. 5uL of 1mM levamisole was added to the pad to anesthetize the worms. The worms were covered with a coverslip and visualized under a fluorescent microscope with GFP filter illumination (395/509 Ex/Em).

#### Histopathology sectioning and analysis for *C. elegans*

Colonization and colonization resistance for the worms was done as previously mentioned. 4 days post colonized worms were then fixed using buffered formalin and then stained with Eosin. The worms were then washed thoroughly with M9 to remove excess stain. Finally, the worms were then mixed with 5% agarose and poured onto a disposable mold. The mold was further filled with 5% agarose. The blocks were then cut into 10um sections and placed on a lysine-coated slide. This slide was imaged using a slide scanner (Motic).

#### Histopathology sectioning and analysis of mice colon

Post euthanasia the complete the gut was exteriorized, and the colon (from cecum to distal colon, without anus) were opened longitudinally. The surface was cleaned and flushed with cold PBS, and the tissue was rolled transversally with the luminal surface inside. The rolls were fixed by using a wooden tooth pick, preserved in formaldehyde for 24 hours, and then placed in 100% ethanol until paraffin embedding. Tissues were embedded in paraffin and sectioned into 4 μm thick slices prior to staining with hematoxylin and eosin. A board-certified veterinary pathologist microscopically examined the slides and the interpretations were based on standard histopathological morphologies. The pathologist, who was blinded to the treatment, compared the treatment sections to the controls. Histopathology was scored on a histomorphological scale from 0 to 3 (normal to severe abnormality) for each of the following: Extent of inflammation, mucosal hyperplasia, crypt architectural distortion, erosion, mononuclear infiltrate, polymorphonuclear and leukocyte infiltrate^9^.

#### Nile Red staining

We performed Nile Red fat staining of post colonized animals as mentioned previously^10^. Briefly, the worms were washed off from NGM Petri dishes with PBS + 0.01% Triton X-100 (PBST) into a 1.5 mL tube, targeting approximately 100 worms per strain. The samples were centrifuged at 25 × g for 1 minute, and the supernatant was reduced to 0.1 mL and the washing process repeated until the supernatant appeared clear. The worms were then fixed with 600 μL of 40% isopropanol for 3 minutes at room temperature. A Nile Red staining solution was prepared by diluting a 0.5 mg/mL stock solution in 40% isopropanol. Following the fixation, the samples were centrifuged again, the supernatant reduced, and 600 μL of the staining solution was added, with an incubation period of 2 hours in the dark. Subsequently, the samples were centrifuged, the supernatant replaced with PBST, and incubated for an additional 30 minutes in the dark. Approximately 14 μL of the worm suspension was then mounted on a microscope slide, covered with a coverslip. The samples were imaged using a Zeiss LSM 980 with Airyscan 2 with a 20× objective and a red (514) filter to visualize the worms.

### Bioinformatics Analysis

#### Phylogenetic tree and distance analysis

The whole genome for the bacteria used in the colonization trial as mentioned previously^2^ and the natural microbiome members^11^ were used. Raw FASTQ files from the two studies were downloaded from the NCBI database, assembled using SPAdes (v3.15.5)^12^, and annotated with Prokka (1.14.6)^13^. The 16S rRNA sequences were then extracted using Barrnap (0.9)^14^. For phylogenetic analysis, essential R libraries such as ’ape’^15^ and ’vegan’^16^ were utilized. The pre-constructed phylogenetic tree was read using read.tree from the ’ape’ library, and cophenetic distances were computed to assess the phylogenetic relationships.

#### GC content analysis

For GC content analysis, a custom R function, calculate-GC-Content, was developed using the ’Biostrings’ package^17^. This function read the sequences from FASTA files and calculated the GC content by aggregating guanine and cytosine nucleotide counts across all sequences, providing a measure of the genomic GC proportion of the bacteria.

#### tBLASTn Analysis

The whole genome for the bacteria used in the colonization trial as mentioned previously^2^ and the natural microbiome members^11^ were used. Raw FASTQ files from the two studies were downloaded from the NCBI database, assembled using SPAdes (v3.15.5)^12^. The FASTA file generated were converted into BLAST database using makeblastdb (2.14.1) tool. Query gene sequences downloaded from NCBI database as mentioned in (Table S2, S3) were made onto a FASTA file and used as query sequence. Finally, we ran tBLASTn (2.14.1) search using the specified query file, saving the results in a TSV format with detailed alignment information. The results were plotted as frequency histogram without an cutoff using R.

#### Random forest analysis of Traitar data

Previously analyzed Traitar data from ^2^ was used for the analysis. For the Random Forest analysis, the ’randomForest’ package^18^ in R was employed, utilizing a dataset preprocessed to select relevant features. The colonization variable was transformed for binary classification (1 for colonized, 0 for non-colonized). A seed value of 123 was set for reproducibility before applying the Random Forest model, which consisted of 500 trees, ensuring robustness in the analysis. The algorithm was tasked with identifying feature importance, ranking variables based on their contribution to predictive accuracy, and helping understand their influence on bacterial colonization across the dataset.

#### Public CDI metagenome dataset analysis

To assess the association of *B. longum* in CDI conditions, an abundance mapping at the strain level was performed using publicly available gut metagenome sequencing data sets collected from research studies as mentioned (Table S4). For all the metagenome reads, quality trimming and adapter clipping were done with Trimmomatic ^19^. Further, the reads were aligned against the human genome to filter out human reads and were assembled with Bowtie2 v2.3.2 ^20^ and SAMtools ^21^. The resulting contigs were classified taxonomically by k-mer analysis using Kraken2 ^22^, with Kraken 2 standard database. Subsequent species abundance estimation was performed using Kraken-tools ^23^ with a threshold set to ignore species with fewer than 10 classified reads.

#### Bacterial community profiling

Microbiota profiling from 16S sequencing was performed using Vsearch ^24^ as previously used. Briefly, Merging and Quality filtering with a minimum cutoff length of 400 and maximum length of 500 were performed for the fastq files using the Vsearch tool. Singleton and chimeric reads (UCHIME) were removed. OTU selection was performed using VSEARCH abundance-based greedy clustering. OTUs were annotated using Custom database made from full-length 16s extracted from the whole genome of the ten bacteria in the consortia.

#### Statistical Analysis

Statistical analyses were performed using GraphPad Prism 9 (GraphPad Software, San Diego, CA, USA) or R. Comparisons between groups were analyzed using either the Mann-Whitney t-test or the t-test with Welch’s correction as mentioned in the figure legends. A p-value of < 0.05 was considered statistically significant.

## Results

### Anaerobic feeding of *C. elegans* L1 larvae with human gut bacteria showed successful colonization

The majority of bacteria in the human gut are obligate anaerobes. In contrast, *C. elegans* is an aerobic organism, which makes feeding with strict anaerobes challenging. To determine whether brief anaerobic exposure could be used during feeding without compromising host viability, we evaluated the survival of *C. elegans* under anaerobic conditions. The nematodes were subjected to an anoxic environment, and periodic assessments of viability indicated that three hours of anaerobic exposure retained viability in more than 50% of nematodes (Supplementary Fig 1).

Capitalizing on these findings, a short-term feeding regimen was adopted in which L1-stage *C. elegans* larvae were fed the previously isolated library of anaerobic human gut bacteria^26^ within an anaerobic chamber for an hour. Subsequently, the larvae were transferred to nematode growth medium (NGM) plates that had been previously seeded with heat-killed *E. coli OP50* as a food source.

For high throughput colonization screen, a 96-well format was utilized to colonize *C. elegans* by previously cultured human gut library ^26^(Supplementary Fig 2a). Given the known defecation rate of *C. elegans* at approximately 45 seconds ^27,28^, we postulated that slow growing bacteria would likely be outcompeted in the nematode’s intestine. Hence, from an initial library of 102 bacterial strains, 19 slow-growing species were excluded, similar to our previous *in vitro* colonization resistance screening against *C. difficile* ^26^ (Table 3).

The colonization experiment resulted in successful colonization of 39 of the 83 bacterial strains fed (Fig 1a). These results collectively suggest that partial anaerobic exposure is conducive to the colonization of *C. elegans* by human gut isolates, thereby providing a novel methodological avenue for microbiome research.

**Figure 1.**
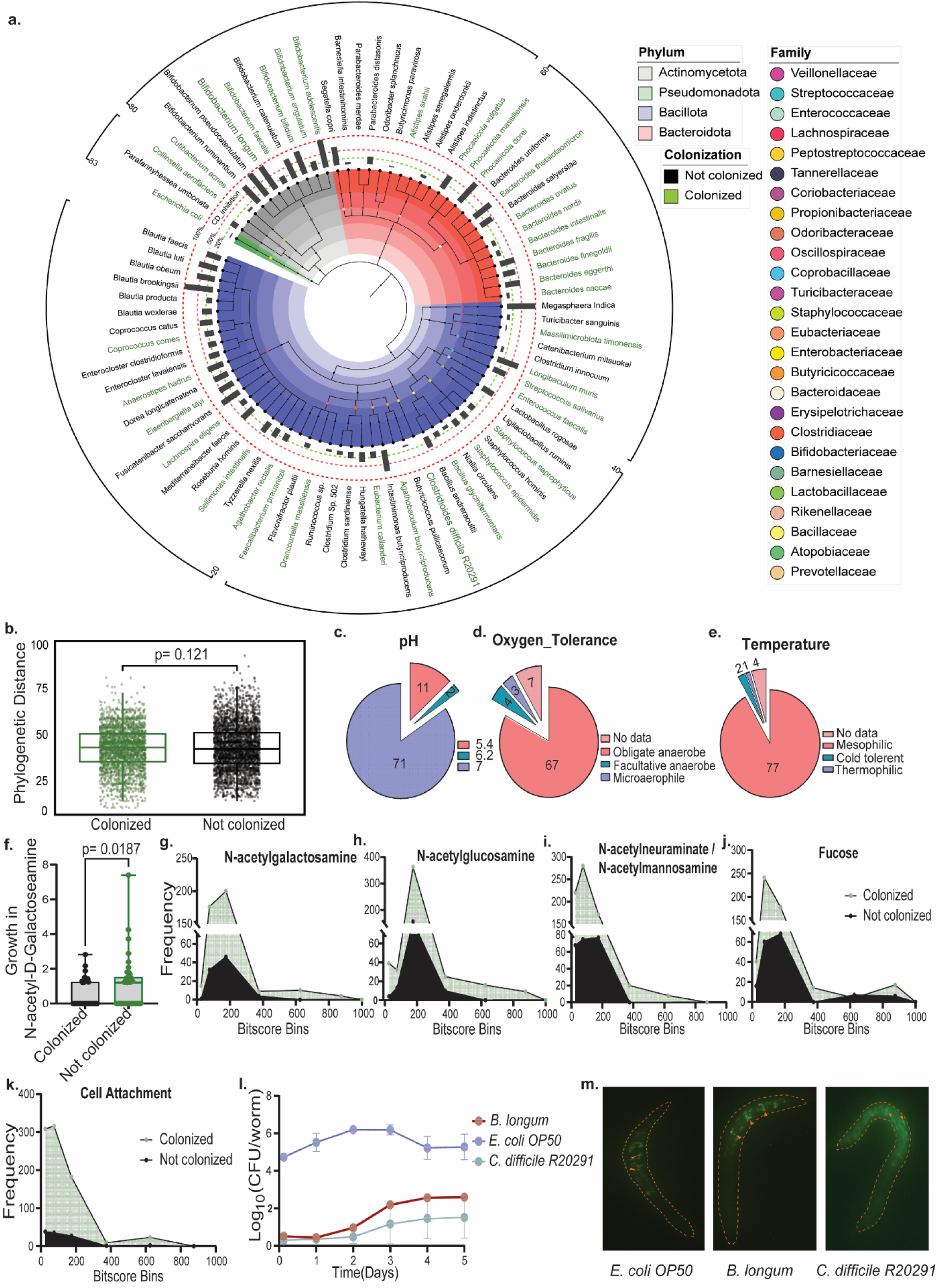
Carbohydrate degradation capability correlates with successful anaerobic bacterial colonization in *C. elegans*. **A.** Displaying a phylogenetic tree derived from 16s rRNA sequence alignment of 82 bacterial species from the human gut and *C. difficile R20291*. Branch colors indicate phylum classification, while circle/text colors represent colonization ability. Surrounding bar graph depicts *C. difficile* inhibition percentages from previous study^26^, with dotted lines at 100%, 50%, and 20% inhibition thresholds. **B.** Median phylogenetic distances between colonized and non-colonized bacteria and natural microbiome members of *C. elegans*^29^, visualized as a scatter plot. Distance calculations are based on pairwise comparisons, with statistical analyses (two-tailed unpaired t-tests) indicated. **C-E.** Pie chart showing the distribution of bacteria based on tolerance to varying pH levels, oxygen, and temperature conditions, informed by predictions for type strains^31^ and supplementary pH data sources^32^. **D.** Comparison of growth between colonized and non-colonized bacteria in minimal media containing N-Acetyl-D-Galactosamine, measured via OD (600nm) and normalized against a water blank. The median line and statistical significance (two-tailed unpaired t-tests) are highlighted. **F-J.** Histogram representing the frequency distribution of bit scores from tblastn for genes involved in the metabolism of N-acetylgalactosamine, N-acetylglucosamine, Fucose, N-acetylneuraminate/N-acetylmannoseamine, and **K.** Cell attachment in the colonized and non-colonized bacterial genome sets, including genome data from natural *C. elegans* microbiome isolates^29^. **L.** Log_10_ CFU/worm graph showing the bacterial counts for *B. longum*, *C. difficile R20291*, and *E. coli OP50* colonized *C. elegans* at various time intervals, with mean ± SEM values from biological triplicates. **M.** Fluorescence microscopy image displaying CFDA-SE stained bacteria in the *C. elegans* intestine post-anaerobic feeding, with arrows indicating colonized bacteria.

### Mucin attachment and carbon utilization ability determine the fate of colonization

To discern the factors determining bacterial colonization in the nematode *C. elegans*, multiple phenotypic and genomic determinants were systematically assessed. Phylogenetic distance-based analyses calculated based on the 16S ribosomal DNA did not reveal a significant relationship between the phylogenetic distribution of colonized and non-colonized bacteria in relation to the natural microbiome members of *C. elegans*^29^ (Fig 1b, Supplementary Fig 2b).

The intestinal environment of *C. elegans* is characteristically slightly acidic^30^, maintained at a cultivation temperature of 25°C, and possesses a degree of anaerobicity due to the size of the gastrointestinal tract. Thus, we examined factors such as pH, temperature, and viability under different oxygen levels to assess colonization dependency. Investigation into these parameters, using phenotypic data from type strains^31,32^ of the bacteria under consideration, did not display a discernible correlation with colonization success (Fig 1c-e & Supplementary Fig 2d-i). Additionally, no correlation was observed between the genomic GC content and bacterial colonization, despite established associations in extremophilic bacterial genomes where carbon sources are limited ^33^(Supplementary Fig 2c).

Carbon utilization capacity is a key determinant of bacterial colonization, a concept extrapolated from dietary dependence in other model organisms. Unlike larger models, such as mice, where gut flora may rely on the host’s diet, *C. elegans* primarily feeds on bacteria, offering mucin as the sole carbon source for colonized bacteria. Analysis of carbon source utilization showed no distinct patterns differentiating colonized from non-colonized bacteria, although colonized strains exhibited a significant preference for N-Acetyl-D-Galactosamine, with no such selectivity observed for other mucin-associated sugars (Fig 1c & Supplementary Fig 5a-g).

Predictive modeling via Random Forest analysis of bacterial genomic traits highlighted spore formation, tartrate utilization, and mucate utilization as key factors determining colonization (Supplementary Fig 4a & Supplementary Fig 3f). The study’s comparison of SCFA production profiles in colonized and uncolonized bacteria showed no significant differences in SCFAs (Supplementary Fig 4b-h).

To probe the genomic basis for mucin utilization, we examined genes implicated in degradation pathways using Biocyc ^34^ predictions (Supplementary Table 1) across both colonized and non-colonized bacterial strains, alongside genomes from *C. elegans’* native gut microbiota. Analysis demonstrated a higher incidence of genes with top Bitscore values for three mucin degradation pathways exclusively in colonized bacteria, barring the fucose pathway (Fig 1g-j & Supplementary Fig 3a-d). Furthermore, an increased occurrence of cell adhesion-related genes was observed in the genomes of these bacteria (Fig 1k & Supplementary Fig 3e).

Collectively, these data underscore the significance of carbon source utilization, with a particular emphasis on sugars inherent to host glycoproteins, in the bacterial colonization of the *C. elegans* intestine.

### Anaerobic bacterial colonization induces differential but temperate responses in *C. elegans* gut integrity, toxicity, and immune dynamics

To study the limitations and extent to which we can use *C. elegans* and the standardized protocol to study colonization and other functional phenotypes, we employed *B. longum*, a known symbiont^35^ with demonstrated inhibitory effects on *C. difficile* in our assays, to test our standardized protocol. Colonization was confirmed through both colony-forming unit (CFU) enumeration periodically post colonization and tagged bacterial imaging inside *C. elegans* intestine. CFU counts confirmed successful colonization by both *B. longum* and *C. difficile*, with observed bacterial colony counts surpassing 1000 CFU per worm over a five-day post-colonization period (Fig 1l). Furthermore, we labeled all three bacteria with Carboxyfluorescein diacetate succinimidyl ester (CFDA-SE) before introducing them to *C. elegans* for colonization. Subsequently, we conducted imaging of the *C. elegans* gut immediately after colonization of *B. longum*, *E. coli OP50*, and C. difficile and successfully visualized them within the intestine (Fig. 1m).

While using *C. elegans* as a model to explore host-microbiome interactions through bacterial feeding is well-established, the prevailing methodology typically employs a continuous feeding regime. This process involves placing *C. elegans* on nematode growth medium plates pre-seeded with specific bacteria, where the nematodes feed and develop until required for further experimental analysis (Supplementary Fig 6a). In the case of *B. longum* and *C. difficile*, we employed a similar approach with a minor adjustment. We seeded both *B. longum* and *C. difficile* onto a NGM plate under anaerobic conditions and subsequently introduced *C. elegans* aerobically to study the host behavioral response post-exposure (Supplementary Fig 6a). Even though this is not a perfect solution to feed anaerobic bacteria, we assume that it will maintain some viability in the cells fed and seek to maintain the freshness of the metabolites produced by the bacteria.

Since we use a different method for colonization (short-term feeding) in *C. elegans* than continuous feeding, we asked whether short-term feeding had a comparable physiological impact to that observed with continuous feeding.

Using the Smurf assay to evaluate gut integrity, we found that short-term feeding with a toxigenic *C. difficile* strain resulted in a higher proportion of leaky gut in *C. elegans* compared to those fed with *E. coli OP50*, although the incidence was lower than with continuous feeding. This pattern held across all tested bacteria, including the pathogenic *C. difficile R20291*, which also caused a notable increase in leaky gut under continuous feeding conditions (Fig 2a-b, Supplementary Fig 6e-f).

**Figure 2.**
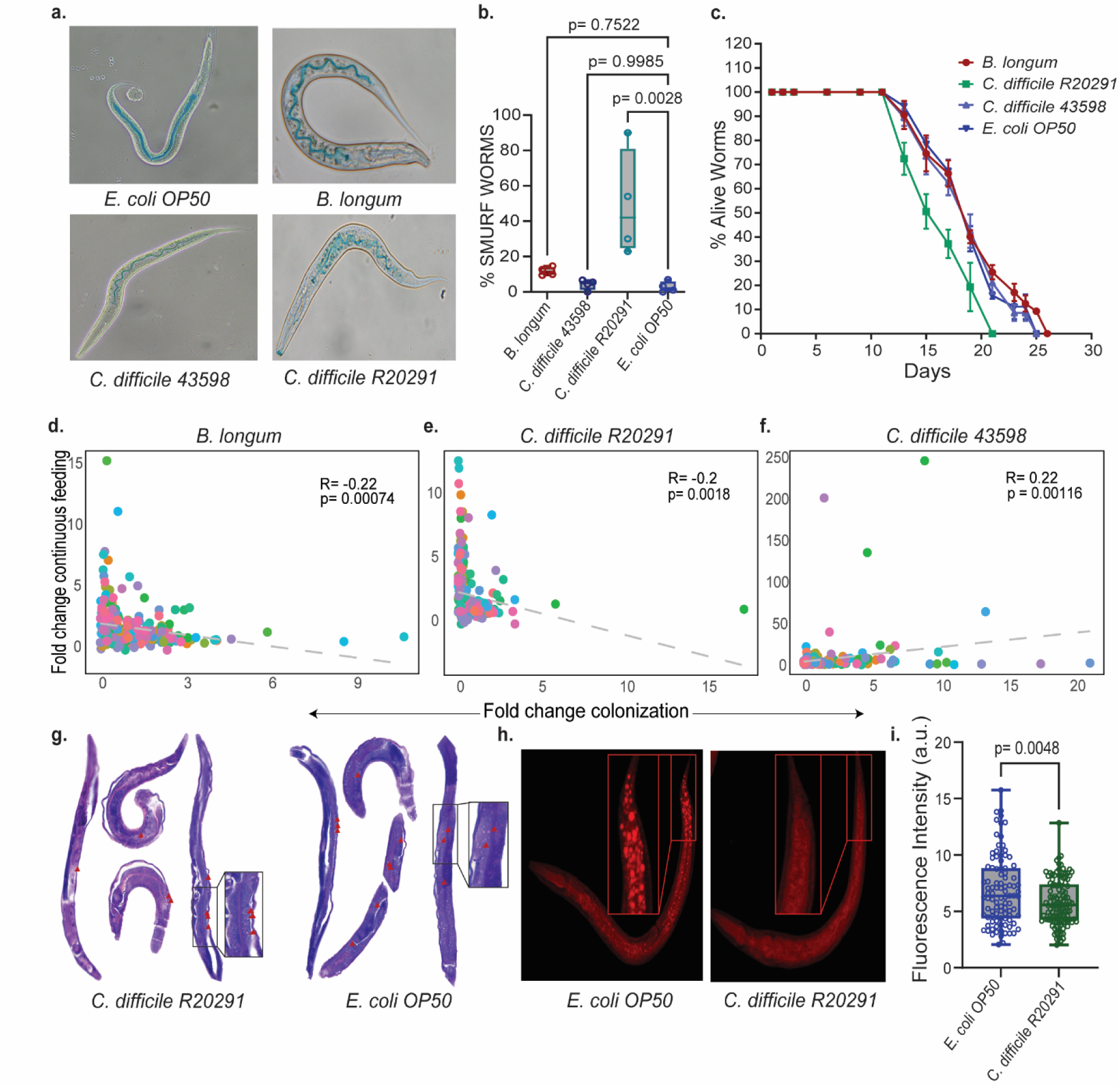
*B. longum* protects *C. elegans* from *C. difficile*-related gut damage and reduced lifespan. **A-B.** Representative images of the *C. elegans* intestine, with a focus on illustrating smurf behavior. Additionally, mean ± SEM values for the percentage of smurf worms are presented for *C. elegans* colonized following the protocol outlined in Figure 1G for *E. coli OP50, B. longum, C. difficile R20291,* and *C. difficile 43598*, with each replicate involving ≥50 C. elegans in biological triplicates. Statistical significance of the observed differences in smurf behavior across the groups is assessed using two-way ANOVA with Dunnett’s correction. **C.** Viability over time shows the mean ± SEM percentage of live worms over time for *C. elegans* colonized using the protocol mentioned in Figure 1G. The worms were colonized with *E. coli OP50, B. longum, C. difficile R20291,* and *C. difficile 43598*. The assay includes is done with ≥50 *C. elegans* per replicate, conducted in biological triplicates. **D-F.** Correlation plots with trendlines indicating the linear relationship between 2^ΔΔCt^ values for continuous feeding, as detailed in Figures 3E-G, and colonization levels as mentioned in Figure S9C-E for include *B. longum, C. difficile R20291,* and *C. difficile 43598*. Each graph also includes the correlation coefficient (R) and the statistical significance of the observed correlations, represented as a p-value. **G.** Microscopic image of eosin stained 10µm thick histopathology section and **H-I.** Fluorescence image of Nile red stained and intensity quantified for *C. elegans* colonized with *C. difficile R20291* and *E. coli OP50* following the protocol outlined in Figure 1G. Mean fluorescence intensity ± SEM is represented as arbitrary units (a.u), with p-value calculated using unpaired students t-tests. Red arrows and zoomed in images highlight the lipid granules.

Lifespan analysis revealed that *C. elegans* subjected to short-term feeding with *C. difficile R20291* experienced lower mortality compared to continuous feeding, albeit higher than those fed with non-toxigenic strains or *B. longum*, which slightly increased longevity (Fig 2c). Median survival under continuous exposure to *C. difficile R20291* was notably reduced, with half the population perishing by 57 hours, underscoring the strain’s pathogenicity; such an effect was not observed with *B. longum* or non-toxigenic *C. difficile 43598* (Supplementary Fig 6b). These findings highlight the potential of short-term feeding in reducing toxicity while maintaining physiological relevance for *C. elegans* research.

The avoidance assay for continuous feeding showed a marked preference for evading the pathogenic *C. difficile R20291*, and unexpectedly, a similar response to the non-toxigenic 43598 strain, suggesting *C. elegans* may recognize both as threats (Supplementary Fig 6c-d). Further testing revealed that while *C. elegans* avoided both *C. difficile R20291 and 43598* supernatants, but the presence of toxin A significantly decreased survival, hinting at a metabolite-driven avoidance mechanism driven by toxin A/B (Supplementary Fig 7a-c). As expected, *B. longum* did not affect the nematodes’ survival or avoidance, akin to the control *E. coli OP50*.

Our investigation into the immune response of *C. elegans* to both continuous and short-term feeding of *C. difficile* strains aimed to uncover any corresponding patterns in immune gene expression. Analyzing the expression of 60 immune-related genes post-exposure to *B. longum*, toxigenic *C. difficile R20291*, and non-toxigenic *C. difficile 43598* revealed a generally weak or negative correlation in gene expression between continuously fed and short-term fed worms, suggesting that short-term feeding induces a more subdued immune response (Fig 2d-f, Supplementary Fig 8, 9 11b-c, & 11e-f).

Specifically, within 12 hours post short-term feeding with *B. longum*, we observed an upregulation in a subset of innate immune genes, notably those within the insulin signaling pathway such as *DAF-16*, without a corresponding increase in its known inhibitor *DAF-2*. These changes align with the longevity benefits noted earlier, suggesting a pivotal role of *DAF-16* in the lifespan extension observed in the nematodes (Fig 2c & Supplementary Fig 11a, d). An increase in *DAF-16* expression was previously noted when *C. elegans* were fed with heat killed *B. longum* and was associated with the observed increase in longevity^36,37^. However, the gene expression changes were transient and appeared to diminish in the later stages of the assay.

Continuous feeding with the pathogenic strain revealed a robust upregulation of the *Scav-2* gene expression, suggesting that *C. elegans* might possess a specialized detection mechanism for anaerobic bacteria, potentially through the *Scav-2* receptor. This is in line with prior documentation of scavenger receptors like *Scav-5* in the immune response to pathogens like *Pseudomonas aeruginosa* and *Salmonella typhimurium*^38^ (Supplementary Fig 11d-f). Additionally, our data showed distinct clustering in gene expression between *B. longum* and *C. difficile R20291*, indicating a differential immune response consistent with the bacteria’s pathogenic profiles (Supplementary Fig 10a-c).

Remarkably, in the nematodes continuously fed with the pathogenic *C. difficile R20291*, a significant increase in the expression of genes associated with the *p38 MAPK* pathway was observed by the 48-hour mark post-infection, mirroring the response pattern to bacterial toxins noted in other models^39,40^ (Supplementary Fig 11e, Supplementary Fig 9). Moreover, the increase in genes related to autophagy in response to the pathogenic *C. difficile* suggested a possible defense mechanism against the bacterial toxins produced (Supplementary Fig 11e). Such a reaction aligns with recent studies in various model organisms, highlighting the conservation of host responses to *C. difficile* toxins^41,42^. Furthermore, the colonization by the non-toxigenic *C. difficile 43598* elicited only a slight alteration in the expression of the *DBL-1* gene shortly after exposure (Supplementary Fig 11b, c).

In summary, our results indicate that while short-term feeding of *C. elegans* leads to successful colonization with lower toxicity, continuous feeding triggers more pronounced immune responses and avoidance behaviors, particularly in reaction to pathogenic strains. This suggests that short-term exposure could offer a more nuanced approach to studying host-pathogen interactions with less stress on the host organism.

### *C. difficile* colonization induces differential lipid accumulation in *C. elegans*

Histopathology plays a pivotal role in infectious disease investigations, providing valuable insights into tissue-level host-pathogen interactions and enabling researchers to visualize and understand the structural and cellular alterations induced by infectious agents. This approach prompted us to explore whether a similar methodology could be applied to *C. elegans*, facilitating an in-depth examination of cellular changes through basic histology staining. After short-term feeding *C. elegans* with *E. coli* and *C. difficile*, we utilized hematoxylin and eosin staining on 10μm-thick sections and observed an intriguing disparity in the distribution of total lipids within *C. elegans* (Fig 2g). Specifically, short-term feeding by the toxigenic strain of *C. difficile R20291* led to a reduction in total lipids, suggesting a potential adaptive strategy employed by the nematodes in response to the reduced lifespan observed upon *C. difficile* colonization, as documented previously in *C. elegans*. To validate the observed difference in lipid accumulation, we performed Nile Red staining to visualize lipids and quantified the staining intensity using microscopy. The result confirmed the observations from the histopathology section, with higher and significant fluorescence intensity shown for *C. elegans* short-term fed with *C. difficile* R20291 (Fig 2h- i). Altogether, our histopathological sectioning approach opens doors to the more pathological exploration in *C. elegans* that has been conducted in microbiome studies.

### Transgenic Analysis Unveils mRNA-Level Regulation of Insulin Signaling Pathway Genes in Colonized *C. elegans*

To further investigate this expression change and its associated physiological response, we utilized a transgene mutant colonization approach. We colonized CF1139 ((pKL78) *daf16*::GFP + *rol-6*(su1006)), a strain with *DAF-16* tagged with a GFP fluorophore, with all four bacteria, and observed an increase in fluorescence intensity at 12, 24 and 48 hours post-colonization, specifically for *B. longum* short-term fed *C. elegans* (Fig 3a, d-g). These findings suggest the possibility of strong RNA-level transcriptional/translational control of *DAF-16* expression. For the toxigenic strain of *C. difficile R20291*, we observed an increase in the expression of only two essential genes involved in antimicrobial defense, *LGG-1* and *GPA-12* (Supplementary Fig 11b). *LGG-1* is part of an autophagy-related pathway^43^, and *GPA-12* is a G protein-coupled receptor in the p38 MAPK pathway^44^. Both genes were significantly upregulated during continuous feeding (Supplementary Fig 11e). This led us to hypothesize a similar level of RNA-level control in their expression. This hypothesis was further confirmed using a transgenic mutant of *LGG-1*. We colonized all four strains in *C. elegans* MAH236 (*lgg-1*p::gfp::*lgg-1p* + *odr-1*p::rfp), where *LGG-1* was tagged with GFP. Upon colonization, we observed an increase in *LGG-1*expression only when the worms were short-term fed with the toxigenic strain of *C. difficile R20291* and *C. difficile 43598* at 12- and 48-hours post-colonization (Fig 3b, h-k). Building on the previous result that demonstrated robust mRNA-level expression control for *DAF-16*, we investigated whether a similar control existed for *DAF-2* gene expression. We used the transgenic mutant QQ202 (cv20 (*daf-2* :: GFP)), where the *DAF-2* gene was tagged with GFP, and short-term feeding it with all four strains. The results confirmed our hypothesis, showing an increase in DAF-2 expression only for the toxigenic strain of *C. difficile R20291* at 12-, 24- and 48-hours post-colonization (Fig 3c, l-o).

**Figure 3.**
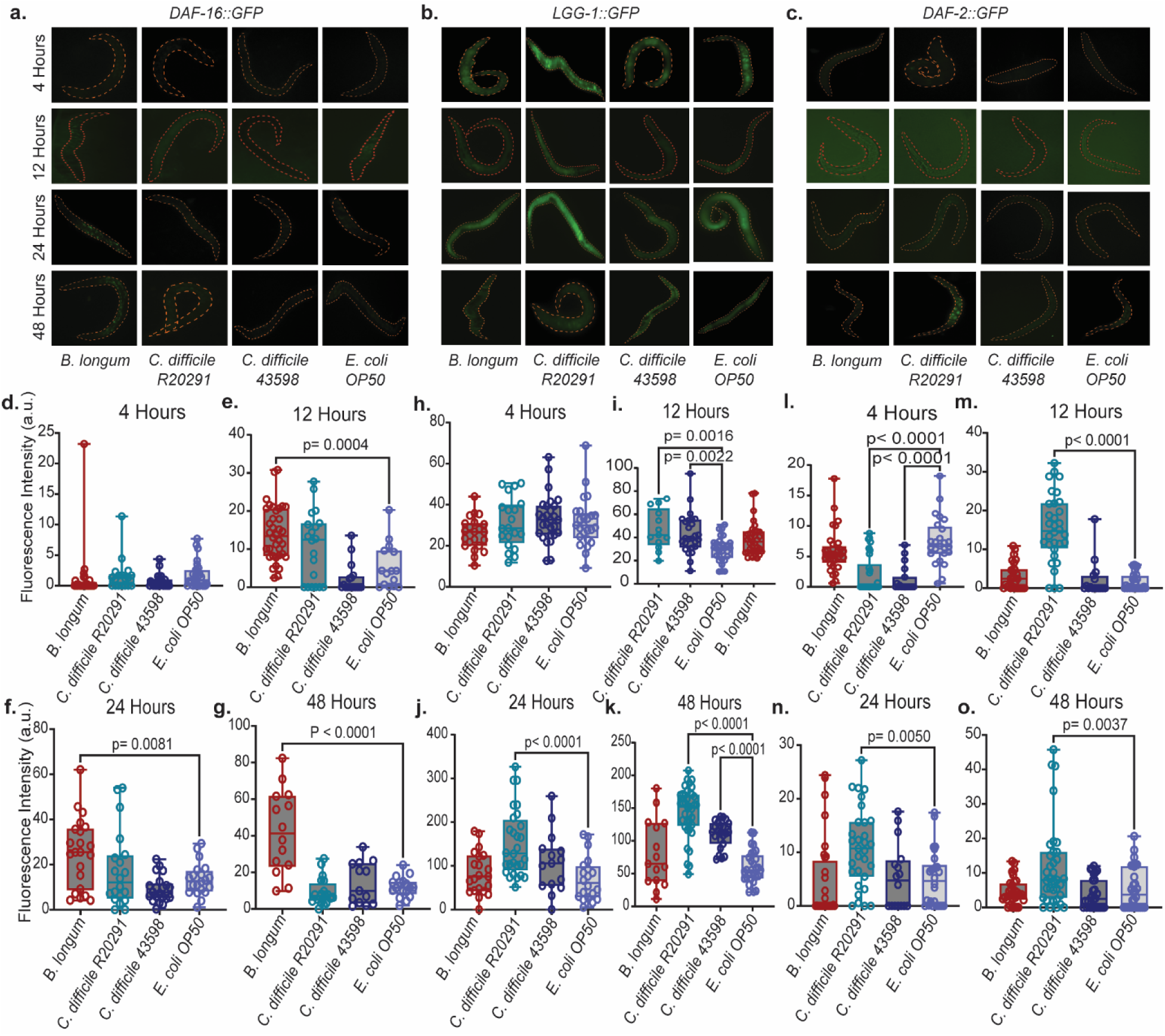
Colonization of transgenic mutants validates toxicity and lifespan responses in *C. elegans*. **A-C.** Microscopic fluorescence images of three different GFP transgene mutants of C. elegans. Each mutant is tagged with GFP for distinct gene markers: **A.** *Daf-16*, represented by CF1139 ((pKL78) *daf16*::GFP + *rol-6*(su1006)); **B.** *lgg-1*, represented by MAH236 (*lgg-1p*::gfp::*lgg-1p* + *odr-1p*::rfp); and **C.** *Daf-2*, represented by QQ202 (cv20 (*daf-2*::gfp)). These mutants were colonized with *B. longum, C. difficile R20291, C. difficile 43598,* and *E. coli OP50* for 4, 12, 24, and 48 hours post colonization, following the protocol outlined in Figure 1G. These images allow for the visualization of GFP expression, providing insights into the localization and regulation of the tagged genes under different colonization conditions. **D-F.** Quantification of the fluorescence intensity observed in the GFP transgene mutants under the various conditions described in **A-C.** The mean fluorescence intensity for each group is expressed in arbitrary units (a.u) and includes the standard error of the mean (SEM). This quantification is conducted for at least 25 worms for every condition, ensuring a robust analysis. Statistical significance of the fluorescence intensity across different conditions and time points is determined using unpaired students t-tests.

Investigations using transgenic *C. elegans* clearly reveal mRNA-level control of insulin signaling genes, with colonizing using short-term feeding amplifying the regulation, particularly in response to *B. longum* and toxigenic *C. difficile* strains.

### *B. longum* pre-colonized worms showed improved gut integrity, rescued lifespan, and colonization resistance against *C. difficile*

We endeavored to standardize a protocol for using *C. elegans* to study colonization resistance against *C. difficile*. Our modified approach involved initiating *C. difficile* colonization a day following the introduction of the top 10 bacteria that exhibited significant inhibition *in vitro* (Fig 1a), with subsequent *C. difficile* quantification using selective media. The enumeration demonstrated a marked decrease in CFU loads for *B. thetaiotaomicron*, *E. tayi*, *B. adolescentis*, *B. eggerthi*, and *B. longum* in comparison to the control *C. difficile* (Fig 4a).

**Figure 4.**
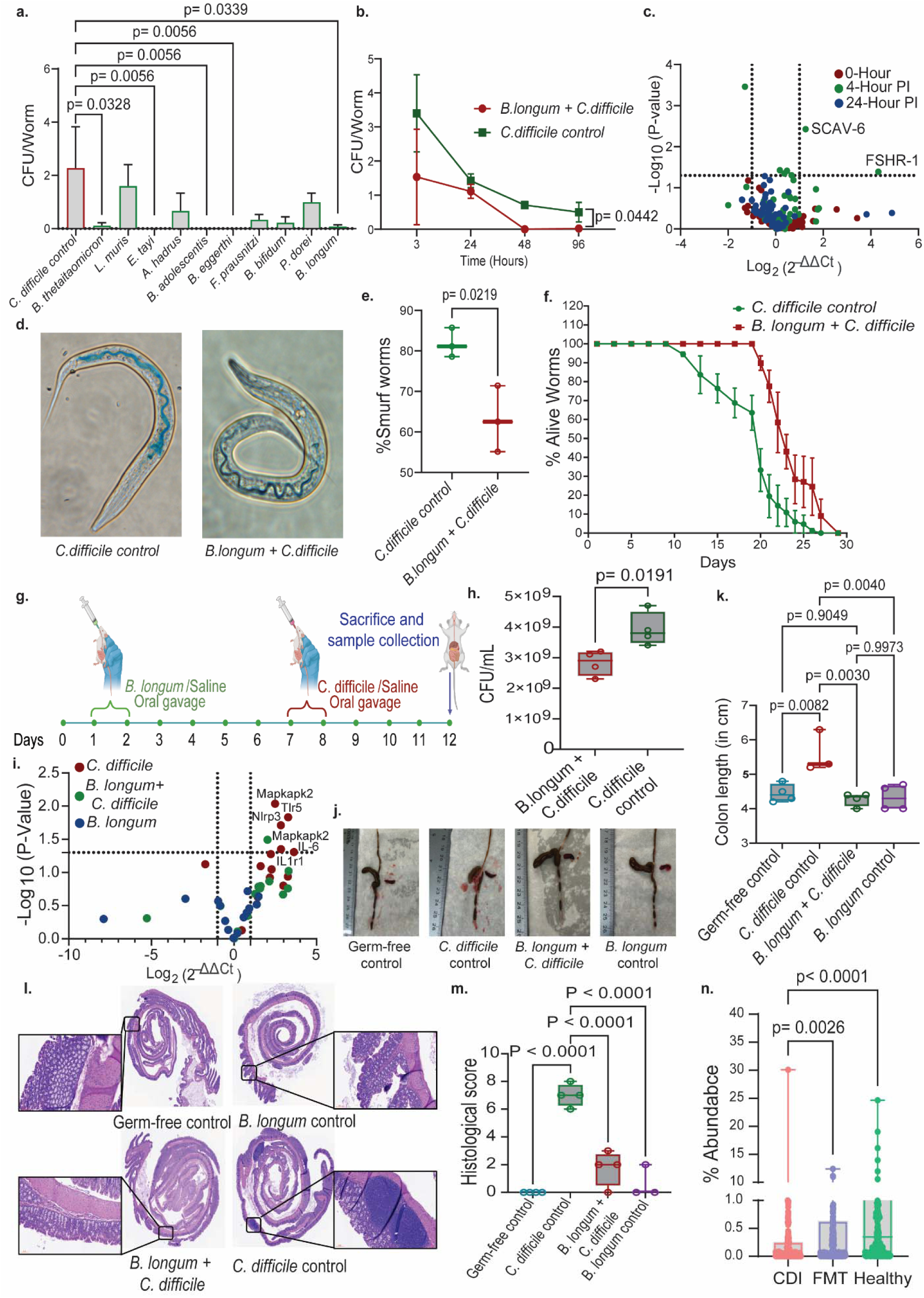
*B. longum* demonstrated effective colonization resistance to *C. difficile* in mice, resulting in a parallel enhancement of intestinal integrity. **A.** Log_10_ CFU/Worm graph showing the counts of *C. difficile* in *C. elegans* that were pre-colonized with *B. longum* or fed control dead *E. coli OP50*, followed by *C. difficile R20291* infection. The data is represented at various time intervals, with mean ± SEM values from biological triplicates. Statistical significance is determined using unpaired students t-tests. **B.** Log_10_ CFU/Worm graph showing the counts of *C. difficile* in *C. elegans* that were pre-colonized with *B. longum* or fed control dead *E. coli OP50*, followed by *C. difficile R20291* infection. The data is represented at various time intervals, with mean ± SEM values from biological triplicates. Statistical significance is determined using unpaired students t-tests. **C.** Volcano Plot displays gene expression data quantified using 62 innate immune-associated genes in *C. elegans* post *C. difficile* feeding for *B. longum* pre colonized worms at 0, 4, and 24 hours. Fold change is calculated using the 2^ΔΔ^Ct method with *E. coli OP50* fed and *C. difficile* infected expression data as control, and P-Value is calculated using unpaired students t-tests. **D-E.** Representative images of *C. elegans* intestine, focusing on the display of smurf behavior, and Mean ± SEM values for the percentage of smurf worms in *C. elegans* colonized with either *B. longum* or fed with *dead E. coli OP50* and subsequently infected with *C. difficile R20291*, 24 hours post-infection. The P-Value is calculated using unpaired students t-tests. **F.** Mean ± SEM percentage of live worms over time for both control groups – those pre-colonized with *B. longum* and those fed with *E. coli OP50*, following infection with *C. difficile R20291*. **G.** Schematic representation of the experimental workflow used to assess colonization resistance in mice treated with *B. longum*. The schematic provides a clear overview of the experimental design, including the treatment and observation timelines. **H.** CFU/mL of *C. difficile R20291* in fecal samples collected from the cecum of mice 4 days post-infection (DPI) after euthanasia. The data compares control mice treated with *C. difficile* only and mice pre-treated with *B. longum*. **I.** Volcano plot for RT-PCR expression data of 14 immune genes associated with *C. difficile* infection in the mice colon. The groups compared include mice treated with *C. difficile* only, mice pre-treated with *B. longum* and then challenged with *C. difficile*, and mice treated with *B. longum* only. Fold changes in gene expression are calculated using the 2^ΔΔCt^ method with expression data from Germfree mice as the control. P-values are calculated using unpaired students t-tests. **J.** Images of the cecum, colon, and spleen from the experimental mice and **K.** Colon length measurement in centimeters for various groups: Germfree (GF) control (n=4), *C. difficile* control (n=4), mice pre-treated with *B. longum* and then infected with *C. difficile* (n=4), and *B. longum* control (n=3). Statistical significance of the differences in colon length across these groups is assessed using two-way ANOVA with Dunnett’s correction. **L-M.** H&E stained histopathological sections of the mice colon for the different groups: Germfree (GF) control, *C. difficile* control, *B. longum* pre-treated and *C. difficile* infection, and *B. longum* control and histopathological score is presented as mean ± SEM values for the sum of scores for different categories, as mentioned in Figure S11B. The statistical significance of observed differences in histological scores across groups is determined using two-way ANOVA with Dunnett’s correction. **N.** *B. longum* abundance analysis illustrating the percentage abundance of *B. longum* during *Clostridioides difficile* infection (CDI) and consecutive fecal microbiota transplantation (FMT) treatment, compared to healthy control individuals. Data is extracted from patient samples, with methodological details provided in the study methods section.

To delve deeper into the mechanisms of colonization resistance, we chose *B. longum* as the investigatory candidate, building on prior knowledge of its physiological impact on the host. Time series CFU enumeration of *C. difficile* revealed inhibition at 96 hours post-infection (Fig 4b). Notable increases in the expression of *SCAV-6* and *FSHR-1* were observed, as opposed to controls treated solely with *C. difficile* (Fig 4c). The absence of significant expression changes in other immune genes indicates that the inhibitory effect of *B. longum* on *C. difficile* may not be entirely immune-dependent. Additionally, we did not detect any significant increase in gene expression for the *C. difficile* only control group (Supplementary Fig 11b).

In exploring the protective effect against *C. difficile* induced pathology, we found that pre-colonization with *B. longum* led to a lower incidence of leaky gut syndrome in *C. elegans*, as evidenced by the smurf assay (Fig 4d-e). Furthermore, pre-colonized worms exhibited a reduced mortality rate (Fig 4f), confirming the model’s effectiveness in mirroring *in vivo* colonization resistance.

Finally, we also investigated the variations in colonization and associated physiology when using different larval stages for *C. difficile* colonization. Notably, we observed that colonization with Larval 1 (L1) led to a significant increase in CFU loads within the intestine, while colonization with Larval 3 (L3) initially exhibited a reduction in colonization, eventually getting stabilized (Supplementary Fig 12a). Interestingly, there were no discernible differences in the percentage of leaky gut observed during colonization at different life stages (Supplementary Fig 12b). Furthermore, we noted an elevated death rate in *C. elegans* colonized at L1 compared to those colonized at L3, likely attributed to the higher colonization levels observed at the L1 stage (Supplementary Fig 12c). These findings highlight the significance of the timing of colonization, as it influences both the extent of colonization and disease severity.

Collectively, these results underscore the suitability of *C. elegans* as a model organism for investigating colonization resistance conferred by anaerobic gut bacteria.

### *B. longum-*associated protective effect is recapitulated in a germ-free mouse model of *C. difficile* pathogenesis

While previous studies have demonstrated *B. longum* mediated colonization resistance *in vivo*, they primarily employed conventional mouse models or different strains of *C. difficile*^45–47^. To bridge this gap in our studies, we sought to replicate our findings from *C. elegans* in a well-established model system of germ-free mice. We colonized GF mice with *B. longum* and allowed stable colonization before gavaging them with *C. difficile* (Fig 4g). Stable colonization was noted by daily CFU counting of the feces (Supplementary Fig 13b). Post *C. difficile* infection subsequent examination of cecal CFU loads revealed a significant but modest reduction in *C. difficile* levels following *B. longum* pre-colonization, compared to the control group without treatment (Fig 4h).

Upon euthanasia, we observed visible differences in cecal and colon architecture such as cecum and spleen size and necrosis features in the control group exposed to *C. difficile*, with a decrease in the *B. longum* pretreatment group (Fig 4j-k). We also measured colon length to investigate whether *B. longum* protects mice from *C. difficile*-induced colon shortening. Unfortunately, we did not observe a significant reduction in colon length compared to germ-free control, but intriguingly, *B. longum*-only colonized germ-free mice showed increased colon length (Fig 4j), possibly due to its ability to produce essential SCFA from dietary fibers as detected (Supplementary Fig 14a-d).

Next, we assessed immune gene expression changes in response to *C. difficile* pathogenesis and *B. longum* pretreatment. We selected 14 different genes (Supplementary Table 3) previously known to be activated during *C. difficile* pathogenesis^48^. We observed a significant increase in the expression of *MAPK* signaling genes, ultimately leading to the upregulation of both *IL-6* and *IL1r1*. However, this elevated expression was reduced or lost significance in *B. longum* pre-colonization (Fig 4i). This implies that pretreatment with *B. longum* significantly diminishes the inflammatory response linked to *C. difficile* pathogenesis and does not trigger gene activation in response to the pathogen. These observations are consistent with the orthologous gene expression patterns observed in *C. elegans*. More importantly we checked for improvement in cell architecture and inflammation from the histopathological slides of the whole colon from all the different treatments mentioned above and scored using the scheme adapted from^49^. We found a higher mucosal, submucosal and transmural inflammation with increased polymorphonuclear infiltrate and moderate increase in mononuclear infiltrate for *C. difficile* control mice. This morphology was very significantly diminished or not seen in the case of *B. longum* pretreated mice (Fig 4l-m, Supplementary Fig 14c). Furthermore, to elucidate the importance of *B. longum* in resisting *C. difficile* colonization, we analyzed human fecal metagenomic data from *C. difficile* infection (CDI) patient’s databases. The results were intriguing and supported our hypothesis that *B. longum* plays a crucial role in colonization resistance against *C. difficile* in humans. *B. longum* abundance was significantly lower in CDI patients compared to healthy fecal controls, and this was further supported by an increase in abundance following fecal microbiota transplantation (FMT) treatment (Fig 4n).

In conclusion, our findings indicate that *B. longum* can inhibit *C. difficile* in mice, corroborating the results obtained from the *C. elegans* trials.

### Differential inhibition patterns against *C. difficile* in multispecies colonization models

In extending our investigations on colonization resistance within *C. elegans*, we employed our model to assess whether multispecies consortia could be established similarly to mono-colonization and their efficacy against *C. difficile* colonization. Selecting 10 bacterial species from the top performers in our in vitro colonization screens *Bifidobacterium bifidum, Faecalibacterium praunitzii, Anaerostipes hadrus, Eisenbergiella tayi, Lactobacillus muris, Bifidobacterium longum, Parabacteroides dorei, Bacteroidetes eggerthi, Bacteroidetes thetaiotaomicron,* and *Bacteroidetes adolescentis* which also colonized in *C. elegans* (Fig 1A), we explored different community configurations by creating ten unique bacterial dropout mixes.

Following a slightly modified version of our established protocol, worms were fed with these bacterial consortia and allowed 24 hours to stabilize before *C. difficile* infection (Fig 5a). Enumeration of *C. difficile* 24 hours post-infection demonstrated varied inhibition patterns across the mixes, with five mixes showing greater than 50% inhibition (Fig 5b). Notably, the absence of *A. hadrus, E. tayi, L. muris, B. longum, P. dorei,* and *B. adolescentis* resulted in reduced colonization resistance. Surprisingly, a consortium containing all ten bacteria did not exhibit effective colonization resistance (Fig 5b). Further analysis of community composition highlighted the dominance of the Bifidobacterium genus in most mixes, implicating its significant role in colonization resistance (Fig 5c).

**Figure 5.**
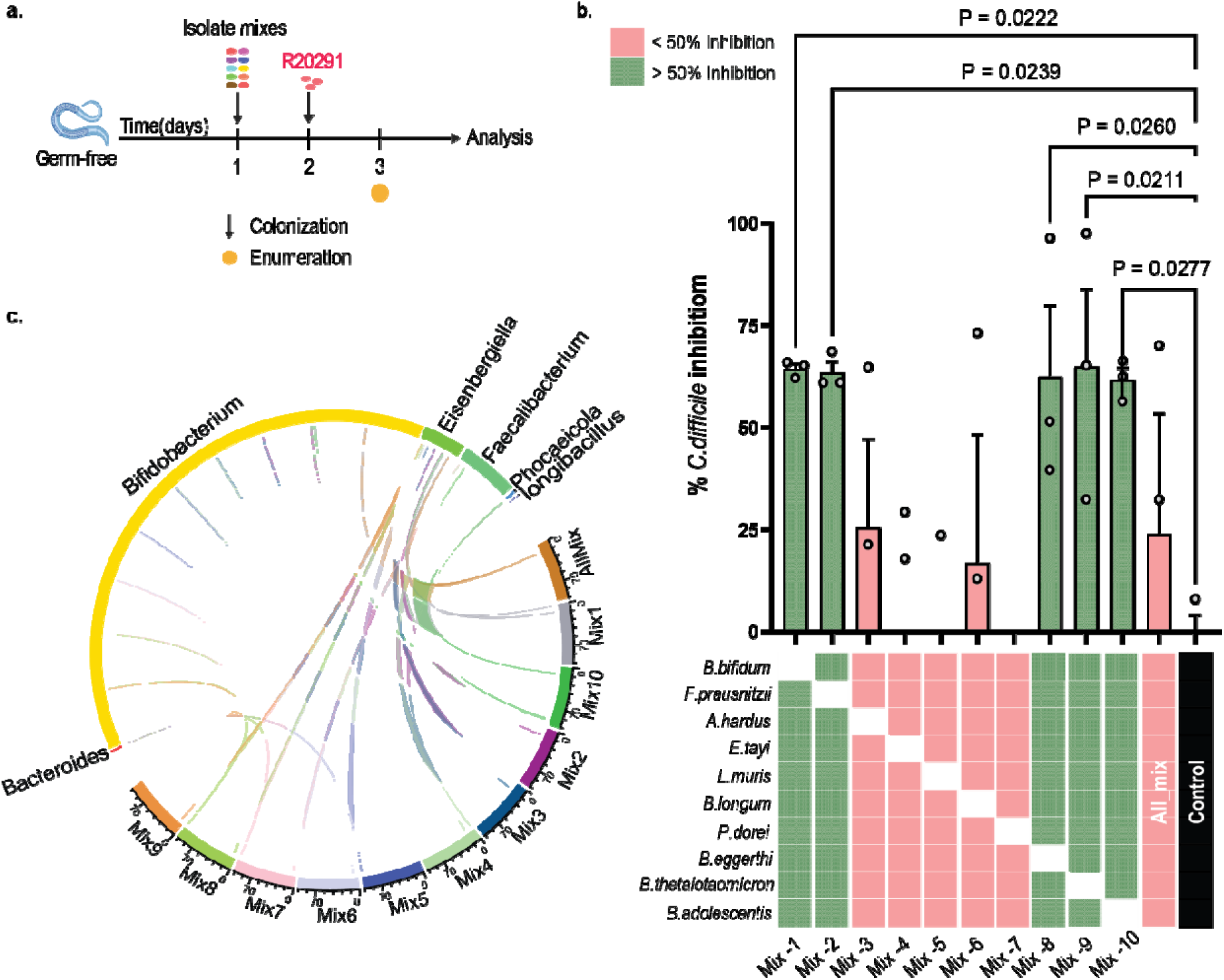
Individual strain dropout from a 10-member consortium reveals mix-dependent inhibition of *Clostridioides difficile* R20291. **A.** Schematic representation of the experimental workflow used to assess colonization resistance against *C. difficile* R20291 using a defined 10-member bacterial consortium and individual bacterial dropout communities (leave-one-out mixes). Each dropout mix was generated by removing one member from the 10-strain consortium, followed by pathogen challenge and quantification of inhibition. **B.** *C. difficile* inhibition across dropout mixes compared to control. Bar plot showing percent inhibition of *C. difficile* R20291, calculated relative to the average CFU of the control condition. Data are shown as mean ± SEM from biological replicates, with individual dots representing replicates. Statistical significance was determined using unpaired Student’s t-tests. X-axis shows matrix representation of inhibition patterns across dropout conditions. Summary plot shows inhibition outcomes across the full set of tested mixes, including the complete consortium (All mix) and individual dropout mixes. Colors classify inhibitory activity based on the threshold criteria: strong inhibition (>50%) (green) and weak/no inhibition (<50%) (pink). **C.** Circos/chord-style representation displaying the relative genus-level abundance profiles of the different dropout communities. Numbers along the track indicate percent abundance (%) of each genus within a given mix, and connecting lines indicate the relative abundance relationships across mixes, highlighting how removal of individual strains reshapes the community abundance structure.

Moreover, there was no significant correlation between individual genus abundance and inhibition patterns, suggesting that higher-order community interactions play a pivotal role in the observed variability of mix-based inhibition.

Collectively, these results affirm the potential of our *C. elegans* model to study multispecies colonization and its impact on colonization resistance. This study underlines the complexity of microbial interactions within poly-microbial communities and their collective influence on pathogen suppression that requires models like *C. elegans* for high-throughput colonization studies.

## Discussion

As the field of microbiome research advances, the application of gut microbiota in the treatment of enteric infections and in the broader context of functional phenotyping and host response screening has become a focal point. This expansion is evidenced by the growth of the culturomics library^50,51^, which poses a challenge in selecting the most promising bacterial species from a vastly enriched pool. Recognizing the limitations of traditional *in vitro* methods, which may not accurately mimic *in vivo* environments, and acknowledging the expensive and laborious nature of common rodent-based *in vivo* models^52–54^, our study advocates for the employment of the nematode *C. elegans*. This organism serves as a versatile, high-throughput model to explore colonization resistance as well as to evaluate functional host responses, offering a more rapid and cost-effective avenue for microbial candidate screening.

*C. elegans* has been utilized as a model organism since the late 1940s to gain insights into host biology. Owing to its microscopic size, the intestine of *C. elegans* cannot be colonized with anaerobic bacteria through simple gavage, which is a technique commonly used in other models. The feasible method to introduce bacteria into its intestine involves exposing the worms to the bacteria of interest, with the expectation that *C. elegans* will ingest them. However, as C. elegans is an aerobic organism, this approach is not effective with obligate anaerobes. Although previous studies have attempted to feed *C. elegans* with anaerobic bacteria^36,37,55,56^, no established method yet exists to effectively colonize the intestine of *C. elegans* with anaerobic bacterial isolates.

For the first time, we have introduced an innovative protocol that enables the colonization of anaerobic human gut bacteria in *C. elegans* without the need for continuous feeding (Fig 1). Our method simulates rodent model gavage by using short-term anaerobic exposure, which has been effective in establishing colonization of a subset of gut microbiota in *C. elegans*. Through genotypic and phenotypic analyses to understand the variability in colonization capacity, we have identified the utilization of mucin-associated sugars like N-acetyl galactosamine and cell attachment factors as critical determinants for colonization (Fig 2). This underscores the importance of mucin as both a nutrient and a supportive element for bacterial proliferation within the host, a finding corroborated by some pathogen-focused studies in *C. elegans*^57–60^. Further investigations revealed that colonization has a more subdued and variable impact on behaviors such as smurf phenotype and lifespan (Fig 3). The altered expression of innate immune genes and a moderated decline in viability indicate that colonization triggers a subtler physiological reaction compared to that of continuous bacterial feeding. The observed discrepancy between the attenuated gene expression response and the pronounced physiological behavioral response prompted us to hypothesize the existence of post-translational controls significantly influencing these changes. Our experiments using transgenic mutants revealed differential protein production for *DAF-2*, *DAF-16*, and *LGG-1* (Fig 4), which was not apparent in RT-PCR results, suggesting the presence of a higher-order post-transcriptional regulatory mechanism that had been previously obscured by continuous exposure and consequent gene overexpression. These findings pave the way for further exploration into novel post-transcriptional regulatory mechanisms or new genetic controls, building upon existing knowledge of *C. elegans*^61,62^.

Our *in vivo* screening for colonization resistance against *C. difficile* included ten bacterial species showed the most inhibition of *C. difficile in vitro*. Our results indicated significant inhibition by five of these bacteria in *C. elegans* when compared to the control. Moreover, we have elaborated on the associated physiology and gene expression on *B. longum* mediated colonization resistance to *C. difficile in vivo*. This was demonstrated by a decreased pathogen burden and reduced adverse health effects in *C. elegans*, as manifested by preserved gut integrity and appropriate immune gene expression. The translational significance of these findings was further affirmed in a germ-free mouse model, reinforcing the validity of *C. elegans* as an effective system for the preclinical assessment of bacterial therapeutics. Our findings also support the mechanism by which *B. longum* inhibits vancomycin-resistant *Enterococcus faecium*, primarily through lumen acidification^63^. This was confirmed using SCFA profiles from the mice’s feces post-colonization.

Finally, our study also advances the understanding of colonization resistance within poly-microbial communities using the *C. elegans* model, highlighting both its utility and the complexity of microbial interactions. Building on our findings with mono-colonized worms, we demonstrated that multispecies consortia could also colonize the *C. elegans* gut effectively and exhibit differential inhibition of *C. difficile* colonization. Notably, five of the dropout mixes achieved greater than 50% inhibition of *C. difficile*, underscoring the importance of specific bacterial members such as *Anaerostipes hadrus*, *Eisenbergiella tayi*, *Lactobacillus muris*, *Bifidobacterium longum*, *Parabacteroides dorei*, and *Bacteroides adolescentis*. Their absence in consortia led to diminished colonization resistance, emphasizing their critical roles. Interestingly, the mix containing all ten bacterial species failed to provide effective colonization resistance, suggesting that higher-order community interactions and potential competitive or antagonistic effects among members may diminish the efficacy of the consortium.

Importantly, the *C. elegans* model has demonstrated its potential as a robust platform for high-throughput preclinical studies of bacterial consortia, capable of unraveling the complex interplay of microbial members that collectively drive colonization resistance. This underscores the translational relevance of our findings, particularly in leveraging microbial consortia as therapeutics against *C. difficile* and other enteric pathogens.

This study has significantly enhanced our understanding of host-microbiota interplay and has also pioneered an innovative and effective method for examining colonization patterns and resistance pathways within the gut microbiota. Although only half of the bacteria assessed exhibited successful colonization, our findings suggest that factors such as sugar metabolism and cell attachment play a pivotal role in these processes. It is postulated that these factor dependencies may vary when the bacteria are introduced as a community or when the medium for C. elegans is altered to better emulate the complex dietary environment of other model organisms. Future research should investigate the combined effects of colonizing and non-colonizing bacteria within a feeder medium that more closely replicates those reported in prior studies. By expanding upon these findings, we can further unravel the intricate dynamics of microbial colonization and its implications for health and disease^64^.

## Data Availability Statement

All 16s amplicon sequencing data generated from this project were deposited in the NCBI SRA database under BioProject . Raw whole-genome sequence data for the strains used were obtained from a previously published study ^65^ under the BioProject <u>PRJNA494608</u>.

## Ethics Statement

All animal procedures were approved by the Institutional Animal Care and Use Committee of South Dakota State University, with prior approval of protocols 19-014A and 19-044A.

## Author Contributions

AA, VTV, PH, SG, and SM performed the experiments, VVP performed the pathological analysis. SG and AR performed NGS and analysis. JS and PK designed the study and provided resources. AA wrote the manuscript with input from other authors.

## Acknowledgments

Computations supporting this project were performed on High-Performance Computing systems managed by Research Computing Group, part of the Division of Technology and Security at South Dakota State University, and the High-Performance Computing Center at the University of Oklahoma.

## Supporting Information

**Figure S1.**
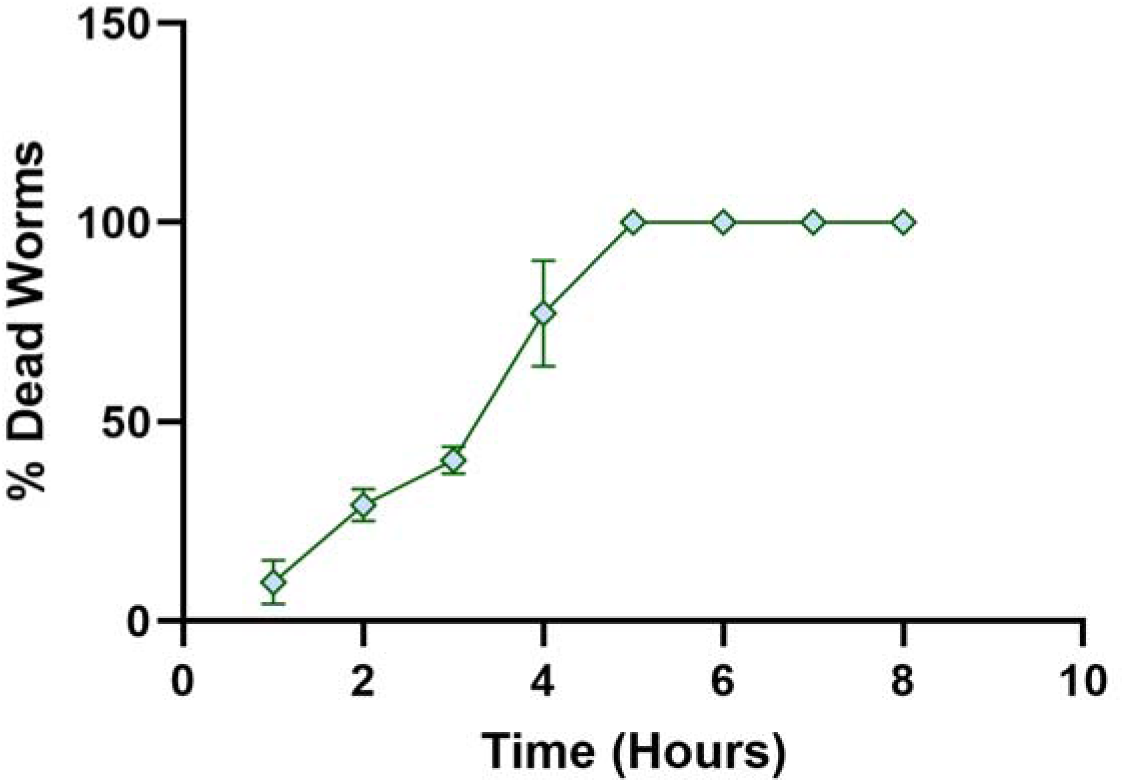
Anaerobicity of *C. elegans*. The figure shows the mean ± SEM percentage of L1 dead worms over time under anaerobic conditions in an anaerobic chamber.

**Figure S2.**
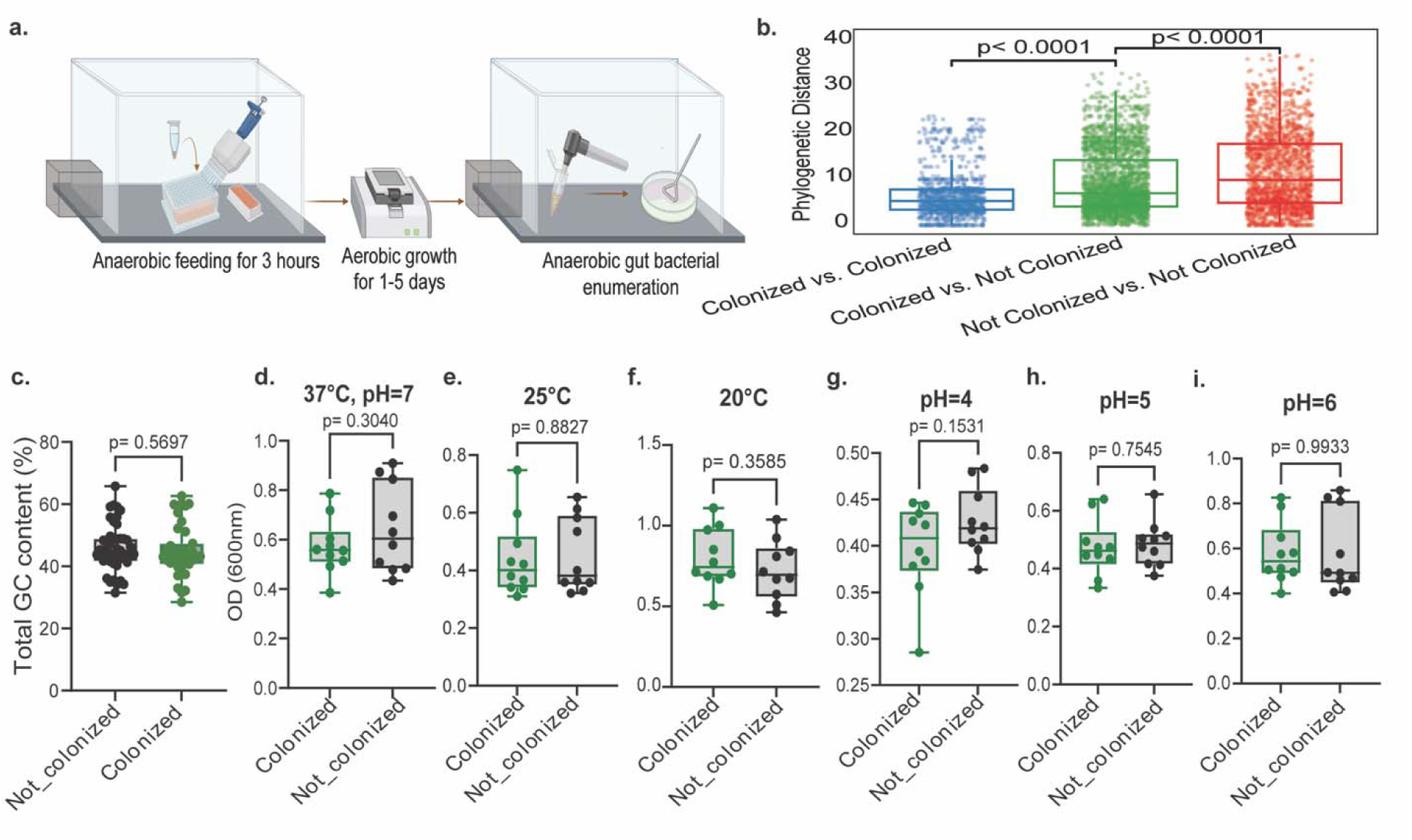
Factors influencing colonization of Human gut anaerobic gut isolates in *C. elegans*, related to Figure 1. **A.** Graphical representation of the methodology used for the colonization of anaerobic human gut isolates in a high-throughput manner. **B.** The figure shows the mean ± SEM values of the phylogenetic distances calculated between and within colonized and non-colonized species. Statistical significance is determined using two-way ANOVA with Dunnett’s correction. **C.** Presented here are the mean ± SEM values for the percentage of GC content identified from the whole genome sequencing of both colonized and non-colonized fractions of bacteria from the human gut microbiome collection. The statistical significance of differences in GC content is calculated using unpaired students t-tests **D-I.** Displayed here are the mean ± SEM OD (600nm) values of 10 blindly selected bacteria from both the colonized and non-colonized fractions of isolates. The growth of these bacteria is assessed post 24 hours at different pH levels (7, 6, 5, and 4) and temperatures (37°C, 25°C, and 20°C). The P-values, calculated using unpaired students t-tests.

**Figure S3.**
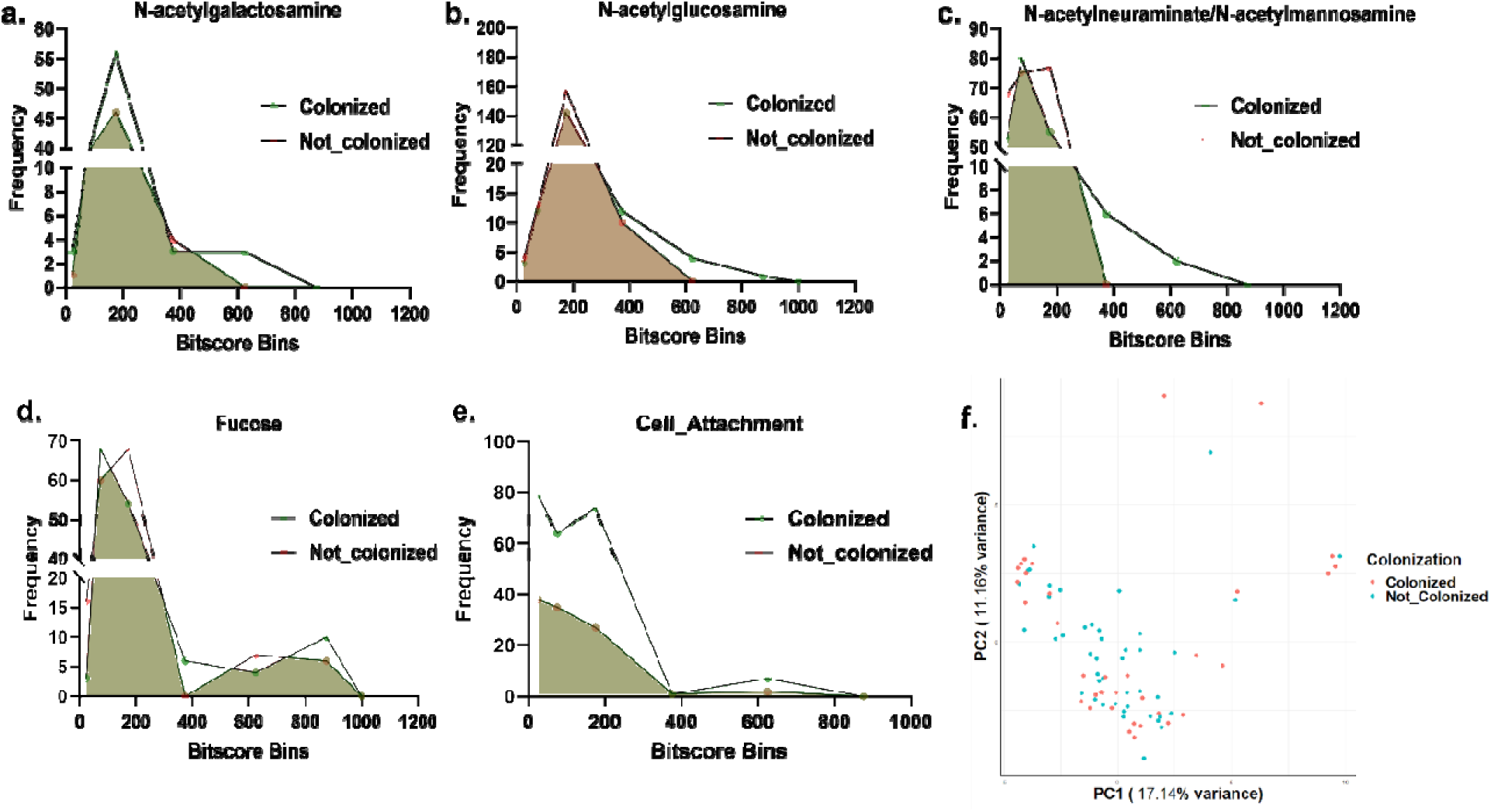
Factors influencing colonization of Human gut anaerobic gut isolates in *C. elegans*, related to Figure 1. **A-D.** The frequency histogram represents the frequency distribution of bit scores from tblastn for genes involved in the metabolism of N-acetylgalactosamine, N-acetylglucosamine, Fucose, and N-acetylneuraminate/N-acetylmannoseamine and **E.** the cell attachment related genes in the colonized and non-colonized bacterial genome sets from the human gut bacterial culture collection used in the colonization trial. **F.** The scatter plot visualizing the results of a Principal Component Analysis (PCA) on Traitar data. The plot displays data points on the first two principal components (PC1 and PC2), which capture the most significant variance in the dataset. Each point represents a bacterial sample, color-coded based on its colonization status.

**Figure S4.**
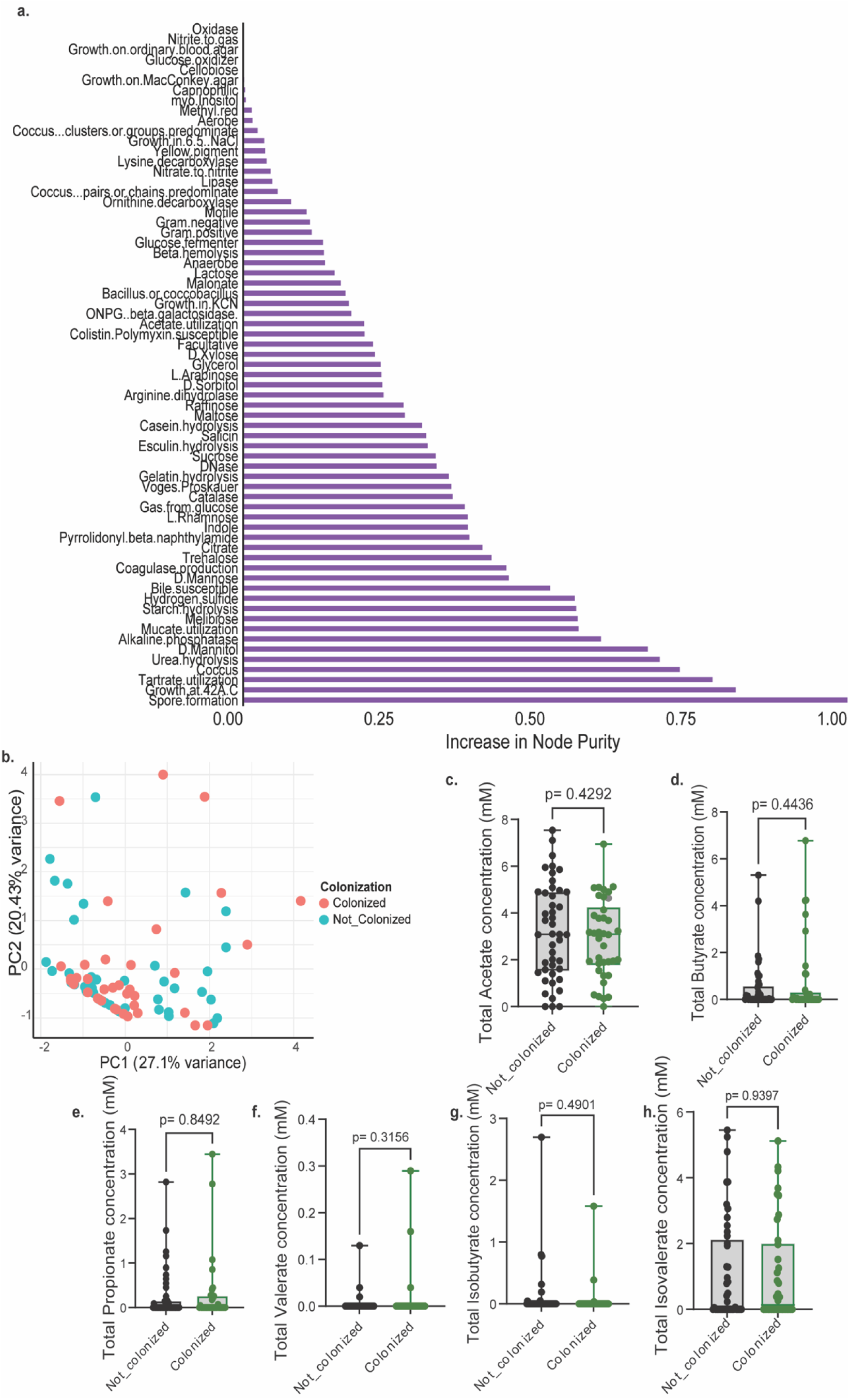
Traitar phenotype measures from colonized and non-colonized, related to Figure 1. **A.** The bar chart displays the results of a Random Forest analysis identifying the importance of different bacterial strains in relation to colonization status. Each bar represents Mean Decrease in Gini Impurity. The chart is designed for ease of interpretation, with strains reordered for clarity and visual impact, highlighting those with the greatest influence on the model’s predictions. **B.** The scatter plot visualizing the results of a Principal Component Analysis (PCA) on Traitar data. The plot displays data points on the first two principal components (PC1 and PC2), which capture the most significant variance in the dataset. Each point represents a bacterial sample, color-coded based on its colonization status. **C-H.** Comparison of **C.** Acetate **D.** Propionate **E.** Isobutyrate **F.** Butyrate **G.** Isovalerate **H.** Valerate concentration between colonized and non-colonized bacteria in mBHI media. The median line and statistical significance (two-tailed unpaired t-tests) are highlighted.

**Figure S5.**
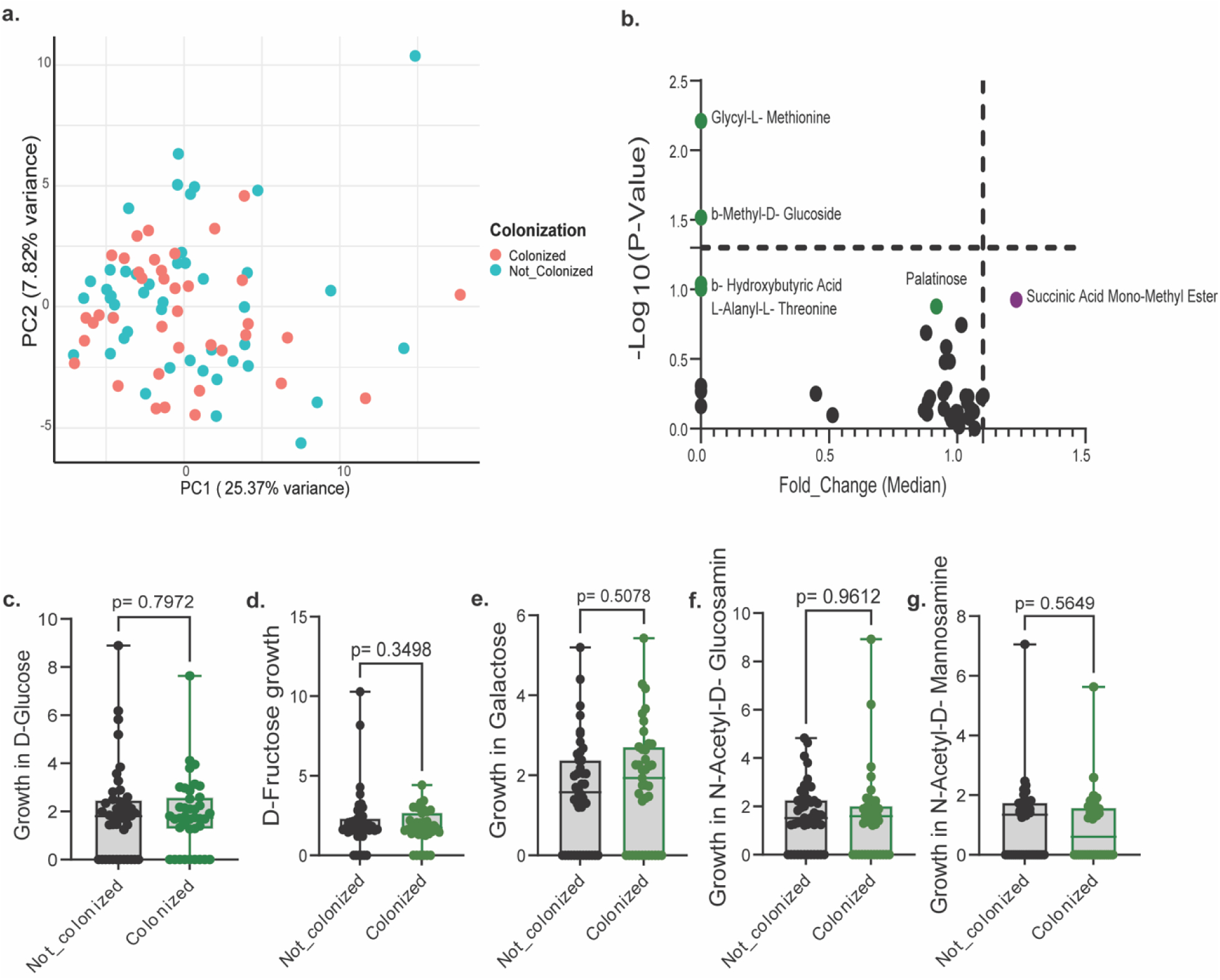
Carbohydrate degradation determination of colonization status, related to Figure 1. **A.** The scatter plot and **B.** Volcano plot visualizing the results of a Principal Component Analysis (PCA) on growth between colonized and non-colonized bacteria in minimal media containing 95 different individual sugars, measured via OD (600nm) and normalized against a water blank. The median line and statistical significance (two-tailed unpaired t-tests) are highlighted. **C-G.** Comparison of growth between colonized and non-colonized bacteria in minimal media containing selected mucin associated sugars **C.** Glucose, **D.**D-Fructose, **E.** Galactose, F. N-Acetyl-D-Glucosamine and **G.** N-Acetyl-D-Mannosamine measured via OD (600nm) and normalized against a water blank. The median line and statistical significance (two-tailed unpaired t-tests) are highlighted.

**Figure S6.**
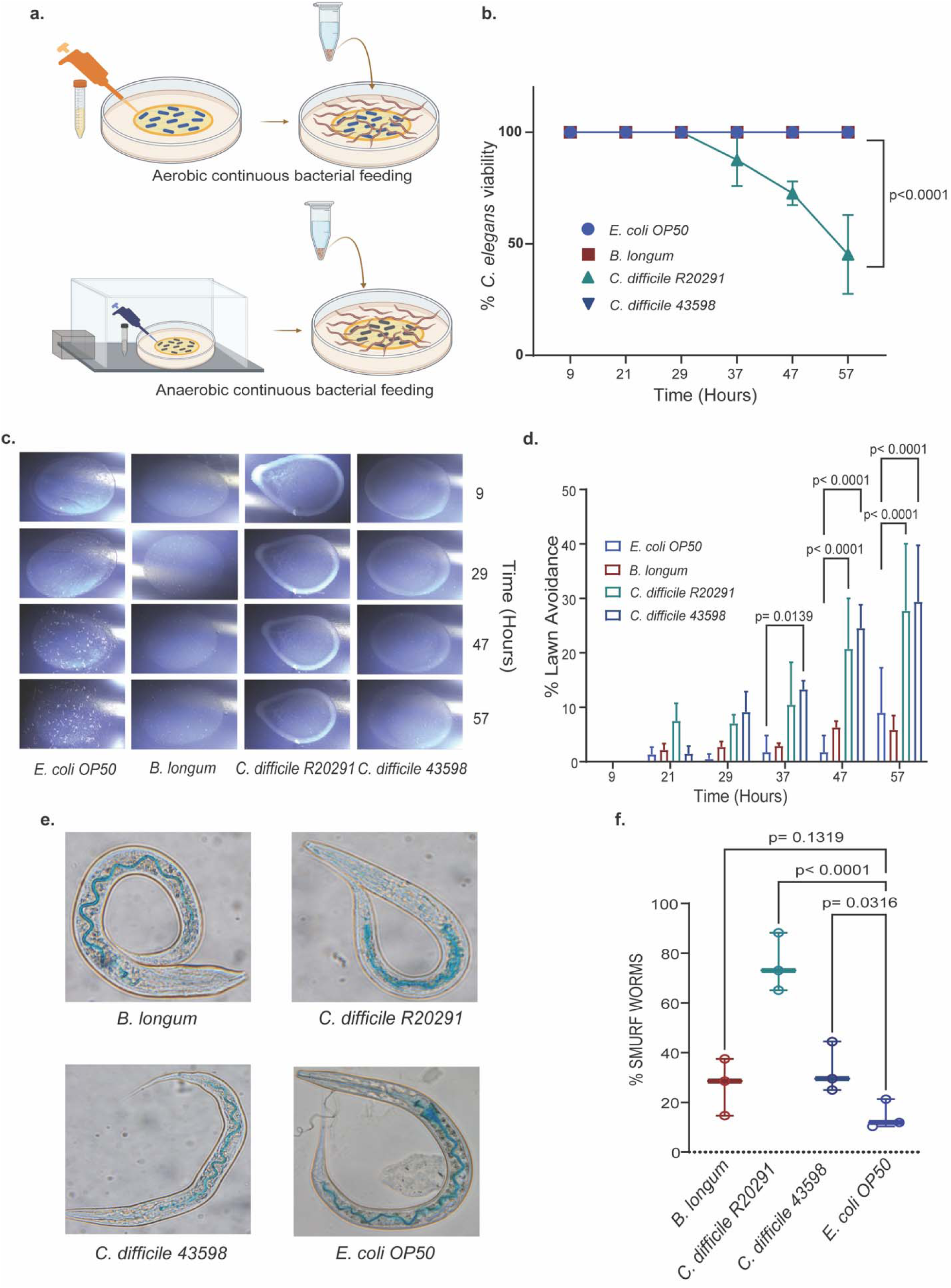
Continuous intake of *B. longum* and *C. difficile* leads to varied avoidance, toxicity, alterations in gut integrity, and immune response in *C. elegans*. **A.** Graphical representation of the methods used for assessing *C. elegans* physiology upon aerobic feeding and a modified version for assessing physiology with anaerobic bacteria. The process involves seeding anaerobic bacteria onto NGM plates within an anaerobic chamber and, after drying, introducing *C. elegans* to these bacteria outside the chamber. **B.** *C. elegans* viability percentage represented as mean ± SEM percentage of *C. elegans* viability after continuous exposure to *E. coli OP50, B. longum, C. difficile R20291*, and *C. difficile 43598*. Measurements are taken at different time points with more than 30 *C. elegans* per replicate in biological triplicates. Statistical significance is determined using two-way ANOVA with Dunnett’s correction. **C.** Avoidance behavior percentage as mean ± SEM percentage of *C. elegans* avoiding the bacterial lawn of *E. coli OP50, B. longum, C. difficile R20291*, and *C. difficile 43598*, measured at the same time points as in **(B).** ≥30 C. elegans per replicate are used in biological triplicates, with statistical significance calculated by two-way ANOVA with Dunnett’s correction. **D.** Representative images of the *C. elegans* intestine, focusing on smurf behavior. Also presented are the mean ± SEM values for the percentage of smurf worms when *C. elegans* are continuously fed with *E. coli OP50, B. longum, C. difficile R20291*, and *C. difficile 43598*, using ≥50 *C. elegans* per replicate in biological triplicates. Statistical significance is assessed using two-way ANOVA with Dunnett’s correction.

**Figure S7.**
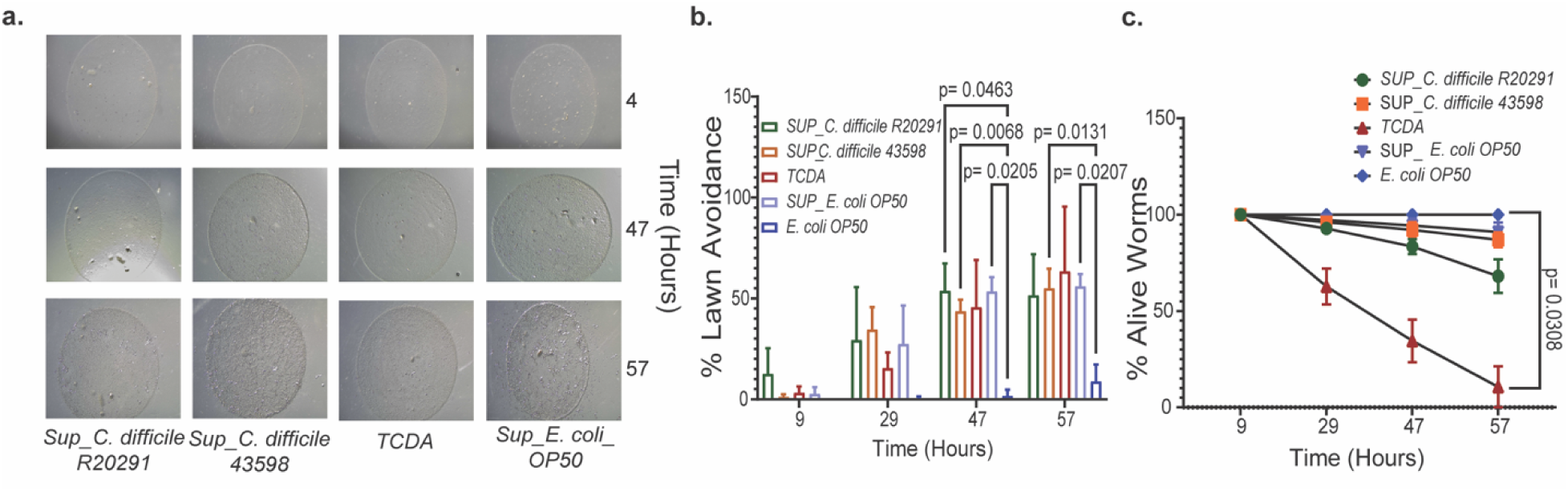
*C. difficile tcdA* toxins decrease viability and avoidance in *C. elegans,* related to Figure 3. **A.** Representative plate images for the bacterial lawn for *C. elegans* fed with dead *E. coli OP50* pellet dissolved in Supernatant from *C. difficile R20291, C. difficile 43598, E. coli OP50,* and 1nM *tcdA* at different time points. **B.** *C. elegans* viability percentage represented as mean ± SEM percentage of *C. elegans* viability after continuous exposure *E. coli OP50* pellet dissolved in Supernatant from *C. difficile R20291, C. difficile 43598, E. coli OP50,* and 1nM *tcdA* at different time points with more than 30 *C. elegans* per replicate in biological triplicates. Statistical significance is determined using two-way ANOVA with Dunnett’s correction. **C.** Avoidance behavior percentage as mean ± SEM percentage of *C. elegans* avoiding the bacterial lawn *E. coli OP50* pellet dissolved in Supernatant from *C. difficile R20291, C. difficile 43598, E. coli OP50,* and 1nM *tcdA* at the same time points as in **(B).** ≥30 C. elegans per replicate are used in biological triplicates, with statistical significance calculated by two-way ANOVA with Dunnett’s correction.

**Figure S8.**
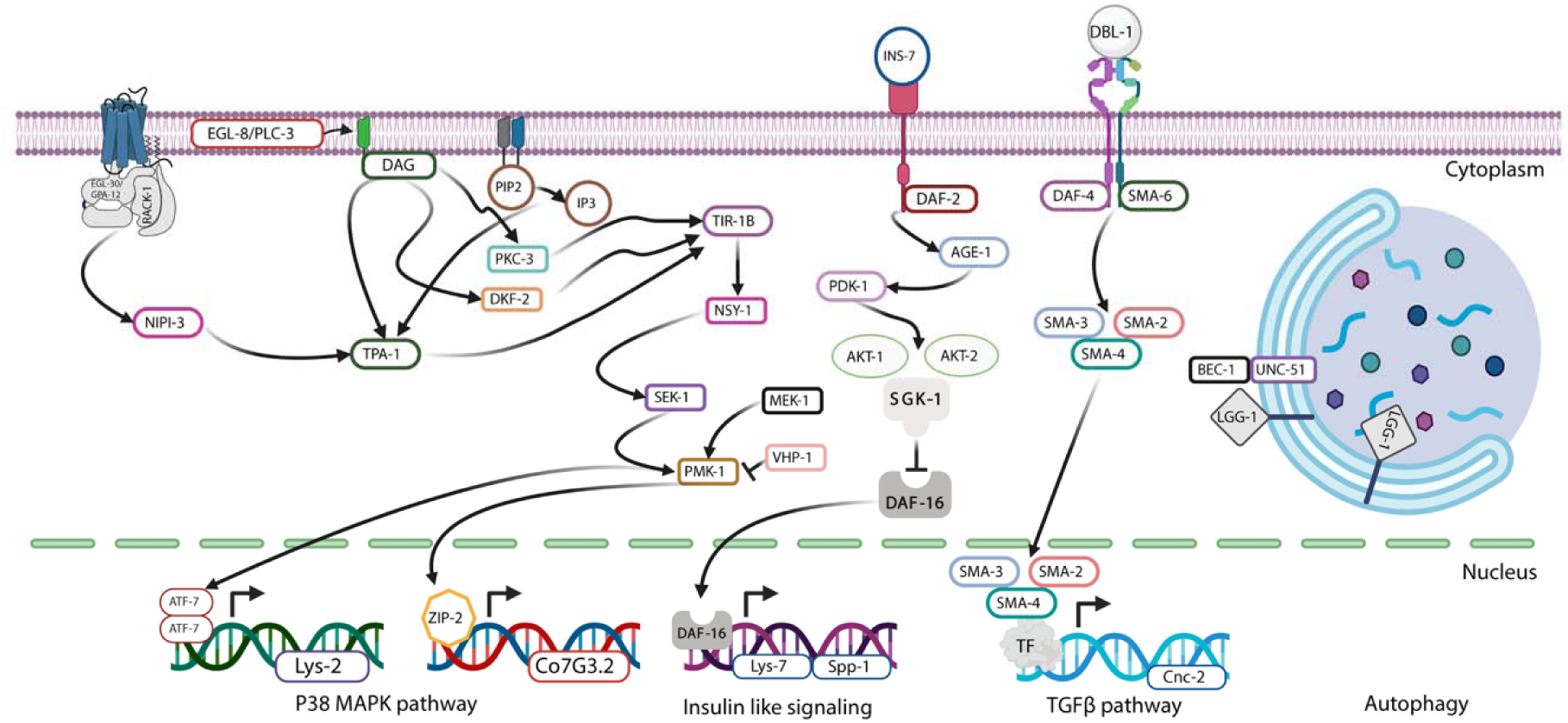
Signaling summary of the selected innate immune genes used for RT-PCR in C. elegans colonization experiment. Related to Figure 2. Created with BioRender.com.

**Figure S9.**
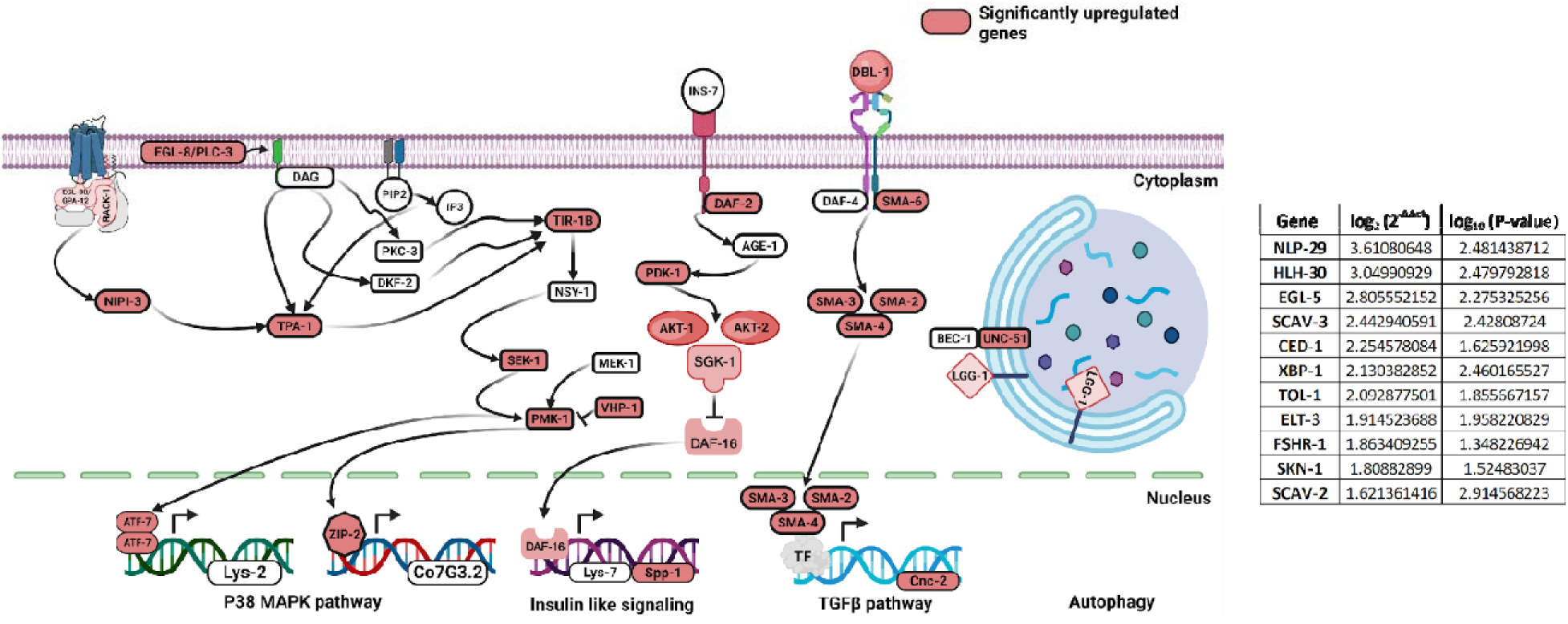
Signaling summary highlighting the genes with red color which are significantly upregulated more than 2-fold and P-value less than 0.05 for only at 48 hours post feeding. The adjacent table indicates the genes present in the pool of 62 innate immune genes which are not part of the signaling pathway mentioned but is significantly upregulated. Created with <u>BioRender.com</u>.

**Figure S10.**
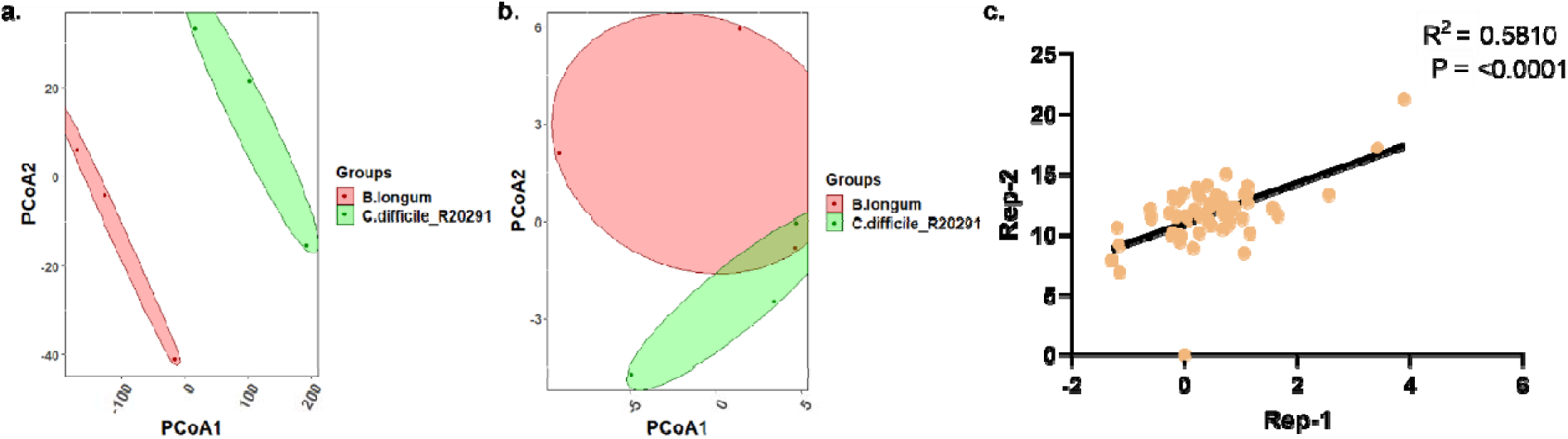
Innate immune gene expression in *C. elegans* shows limited global segregation across bacterial feeding conditions. **A.** The scatter plot visualizing the results of a Principal Component Analysis (PCA) on gene expression data quantified for 62 innate immune-associated genes in *C. elegans* post continuous feeding with *E. coli OP50, B. longum, C. difficile R20291*, and *C. difficile 43598* at 4, 12, 24, and 48 hours as mentioned in Figure 3E-G. **B.** The scatter plot visualizing the results of a Principal Component Analysis (PCA) on gene expression data quantified for 62 innate immune-associated genes in *C. elegans* post continuous feeding with *E. coli OP50, B. longum, C. difficile R20291*, and *C. difficile 43598* at 4, 12, 24, and 48 hours altogether as mentioned in Figure 4C-E. **C.** Replicate gene expression data represented a correlation plot.

**Figure S11.**
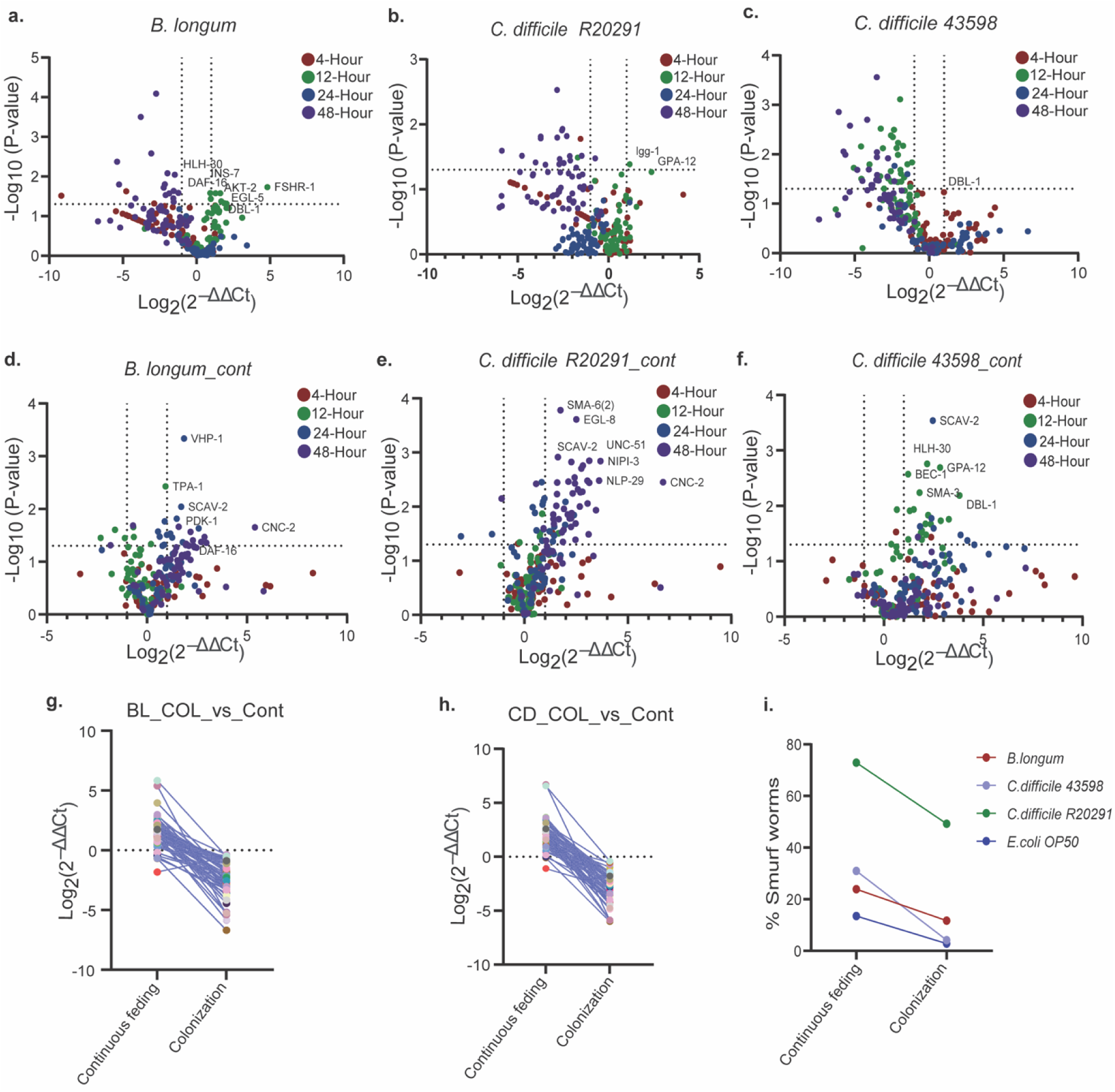
Innate immune gene induction differs between transient colonization and continuous feeding in *C. elegans*. **A-C.** Volcano plots for gene expression data quantified for 62 innate immune-associated genes in C. elegans colonized with *E. coli OP50,* **A.** *B. longum,* **B.** *C. difficile R20291*, and **C.** *C. difficile 43598* at 4, 12, 24, and 48 hours. The fold change is calculated using the 2^ΔΔCt^ method with *E. coli OP50* colonized expression data as the control at the respective time point. P-Values are calculated using unpaired students t-tests. **D-F.** Volcano plots for gene expression data quantified for 62 innate immune-associated genes in C. elegans post continuous feeding with *E. coli OP50,* **D.** *B. longum,* **E.** *C. difficile R20291*, and **F.** *C. difficile 43598* at 4, 12, 24, and 48 hours. The fold change is calculated using the 2^ΔΔCt^ method with *E. coli OP50* continuously fed expression data as the control at the respective time point. P-Values are calculated using unpaired students t-tests. **G-H.** Mean values of gene expression data shown for *C. elegans* colonized and continuously fed with **G.** *B. longum and* **H.** *C. difficile R20291,* as shown in Figure 3E-F and S9C-D. **I.** Mean values for the percentage of smurf worms shown for *C. elegans* colonized and continuously fed with *E. coli OP50, B. longum, C. difficile R20291,* and *C. difficile 43598*, as mentioned in Figures 3D and 4A , showing the difference.

**Figure S12.**
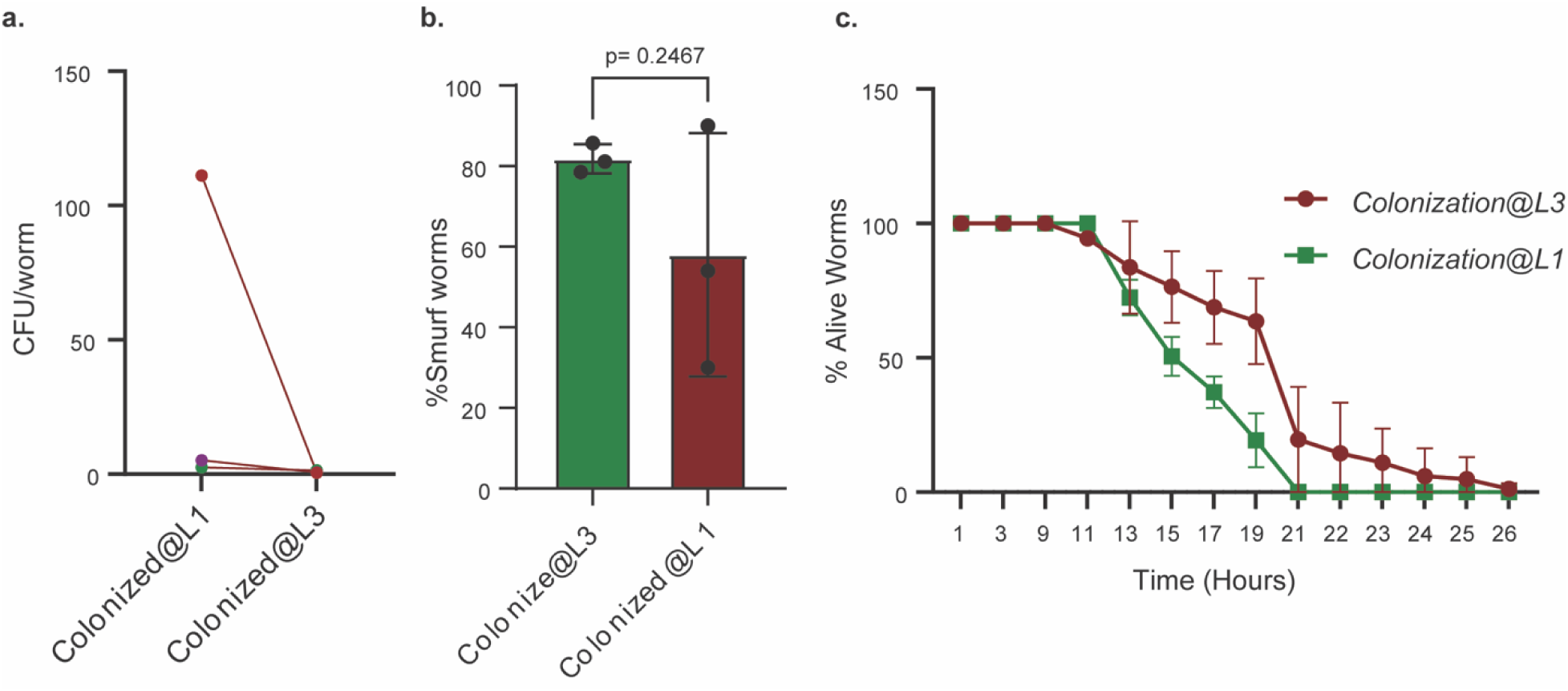
*C. elegans* performs differentially in colonization and disease physiology when colonized at different life stages. **A.** CFU/worm graph showing the bacterial counts for *C. difficile R20291* colonized *C. elegans* at various time intervals for colonized in L1 worms as mentioned in Figure 1H and L3 worm as mentioned 2B. **B.** Mean ± SEM values for the percentage of smurf worms shown for *C. elegans* colonized with C. *difficile R20291* at L3, as mentioned in Figures 2D and L1, mentioned in 4A showing the difference. **C.** Mean ± SEM percentage of alive worms for *C. difficile R20291* colonized *C. elegans* at L1 worms as mentioned in Figure 4B and L3 worm as mentioned 2E.

**Figure S13.**
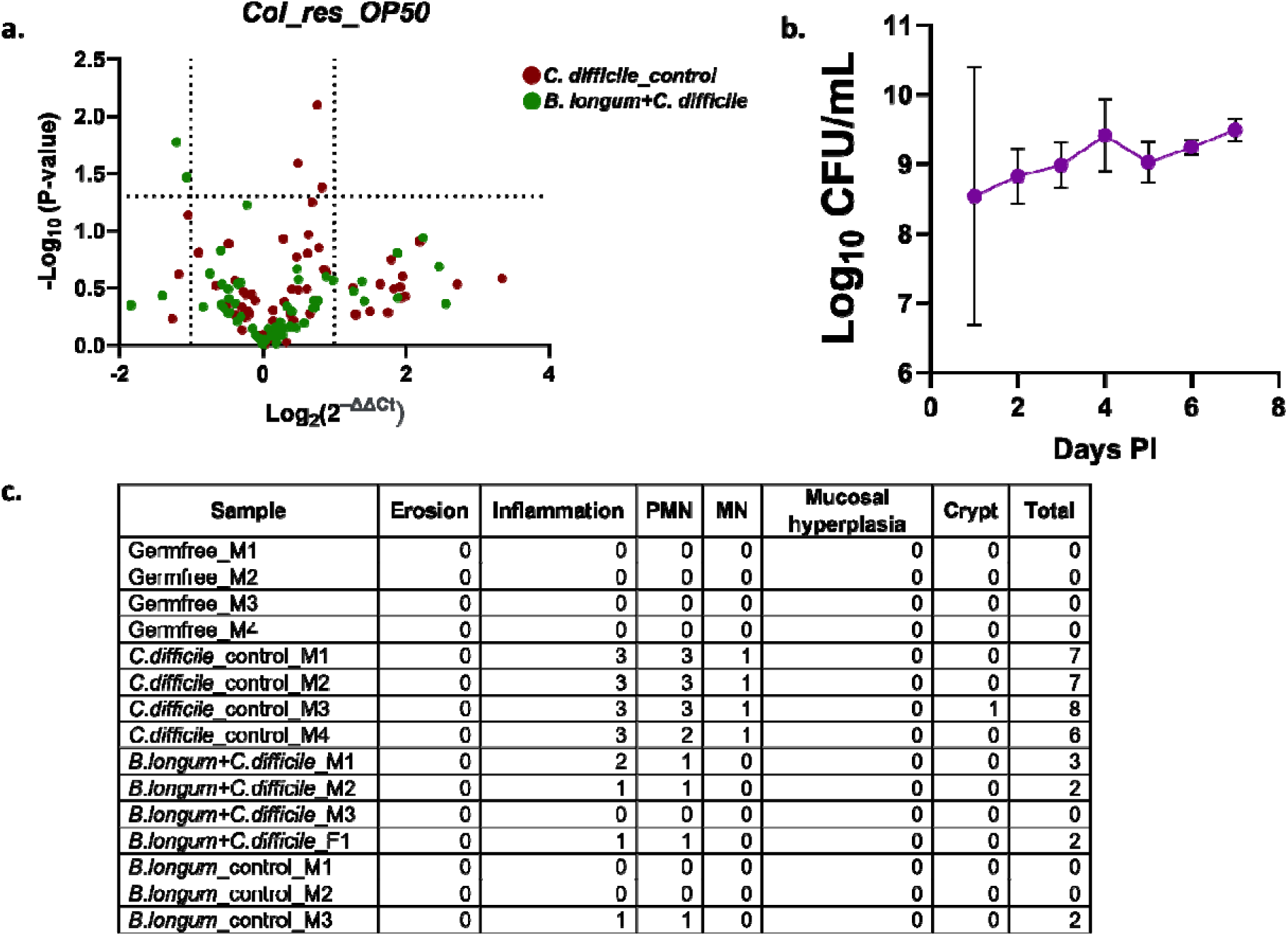
*B. longum* establishes stable colonization in mice with limited host transcriptional shifts in the *C. elegans* model. **A. RT–qPCR analysis of immune-associated gene expression in the** *C. elegans* **colonization resistance model.** Expression of a targeted innate immune gene panel was quantified following **OP50 control feeding**, *C. difficile* **colonization**, or *B. longum* **pre-colonization, followed by** *C. difficile* **challenge.** Overall, gene expression changes were **modest,** with **no major transcriptional differences** observed between Cd-only and BL+Cd challenge groups. Fold change was calculated using the **2^–**ΔΔ**Ct method**, normalized to OP50 controls. Statistical significance was determined using **unpaired Student’s t-tests.** **B.** *B. longum* **colonization in mice.** CFU quantification demonstrates **successful establishment and persistence** of *B. longum* following colonization. Data are shown as **mean ± SEM.** **C. Histopathology scoring of mouse colon tissue.** Colon sections were scored for pathology-associated features (as defined in Methods), and **composite histopathology scores** are presented.

**Figure S14.**
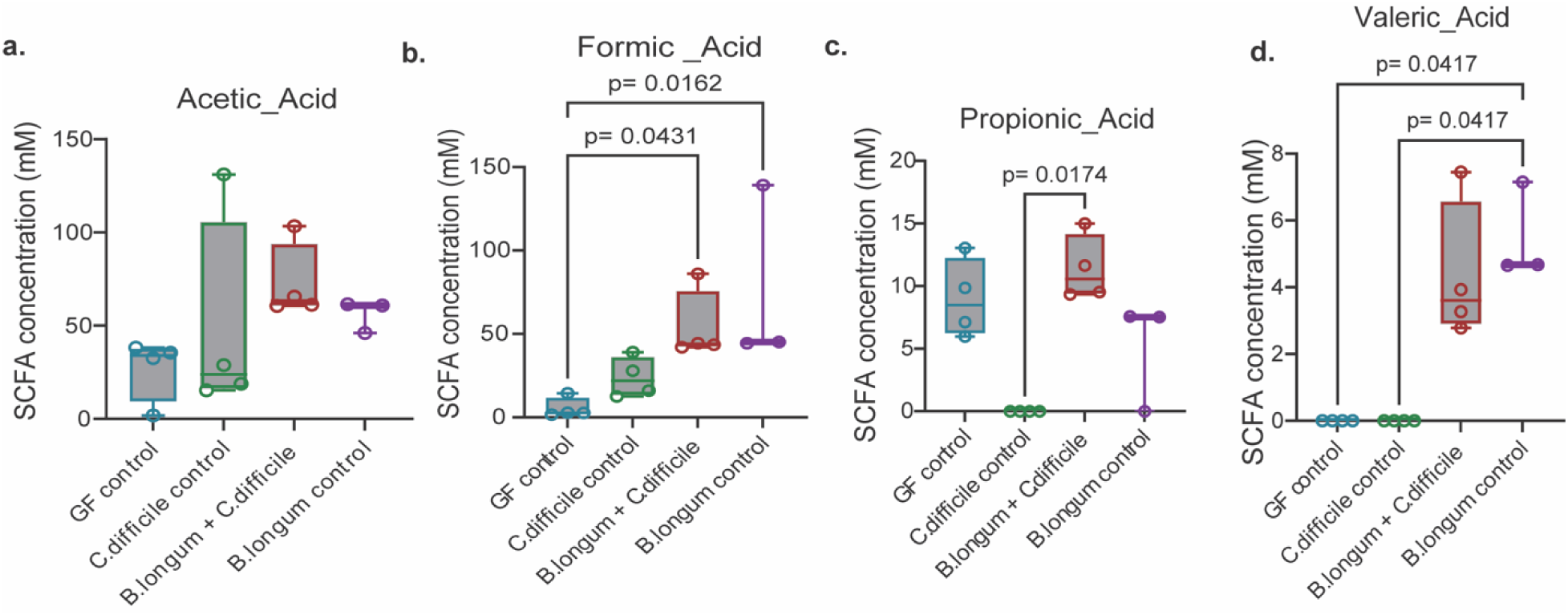
*B. longum* modulates short-chain fatty acid profiles during *C. difficile* infection. **A-D.** Concentration of **A.** Acetic acid, **B.** Formic acid, **C.** Propionic acid, and **D.** Valeric acid identified from the cecal sample of mice colonized with Germfree (GF) control (n=4), *C. difficile* control (n=4), mice pre-treated with *B. longum* and then infected with *C. difficile* (n=4), and *B. longum* control (n=3). Statistical significance of the differences in colon length across these groups is assessed using two-way ANOVA with Dunnett’s correction.

